# Tracking maternal proteins uncovers a central role for the residual body in organelle recycling during *Toxoplasma gondii* replication

**DOI:** 10.1101/2025.08.15.670481

**Authors:** Julia von Knoerzer-Suckow, Parnian Sazegar, Javier Periz, Simon Gras, Markus Meissner

**Author notes:** authors contributed equally to this work (Julia von Knoerzer-Suckow, Parnian Sazegar).

## Abstract

*Toxoplasma gondii* replicates through endodyogeny, an unconventional form of internal budding in which two daughter cells are assembled within a single mother cell. During this process, daughter cells must acquire a full complement of organelles, which may be inherited from the mother, formed *de novo*, or assembled through a combination of both mechanisms. To date, the fate of maternal components during replication remains poorly understood. We previously showed that F-actin–driven dynamics generate an intravacuolar network, associated with the residual body (RB) and facilitates the recycling of microneme proteins. However, the inheritance and recycling of other organelles have not been systematically analysed.

To address this, we employed a dual HaloTag-based pulse-chase fluorescence labelling strategy to distinguish between de novo-synthesized and maternally inherited proteins in replicating tachyzoites. This approach reveals three distinct modes of organelle inheritance: (1) inheritance of intact maternal organelles (e.g., rhoptries and micronemes), (2) expansion and division of pre-existing maternal organelles with incorporation of newly synthesized components (e.g., Golgi apparatus and apicoplast), and (3) degradation of maternal structures followed by de novo assembly (e.g., inner membrane complex). Furthermore, we identify Myosin F (MyoF) as an important factor in the redistribution of maternal micronemes and rhoptries. In the absence of MyoF, these organelles accumulate in the RB, while restoration of MyoF permits accumulated maternal micronemes to redistribute to daughter parasites. Together, these findings support a role for the RB as a dynamic compartment in the recycling and redistribution of maternal organelles.

## Introduction

*Toxoplasma gondii* tachyzoites replicate through a specialized form of internal budding known as endodyogeny, wherein two daughter cells (DCs) are assembled within the cytoplasm of the mother cell (Francia and Striepen, 2014; Hu et al., 2002). This unique replication mechanism demands precise orchestration of organelle inheritance and spatial organization within the parasitophorous vacuole (PV). Apicomplexan parasites, including *T. gondii*, exhibit a highly polarized cellular architecture defined by specialized secretory organelles—micronemes, rhoptries, and dense granules—essential for host cell invasion and intracellular survival. These are accompanied by canonical organelles such as the Golgi apparatus, endoplasmic reticulum (ER), a single mitochondrion, and the apicoplast, a non-photosynthetic plastid. Notably, unlike most eukaryotes, *T. gondii* is enveloped by a pellicle consisting of the plasma membrane and an underlying inner membrane complex (IMC) (Ouologuem and Roos, 2014), a structure composed of flattened vesicles tightly associated with the subpellicular cytoskeleton (Harding and Meissner, 2014).

As DCs emerge during endodyogeny, they progressively encapsulate nearly all maternal cytoplasmic contents. While the sequence of organelle acquisition has been extensively characterized (Nishi et al., 2008), the extent to which maternal organelles and proteins are recycled remains unclear. Historically viewed as a passive remnant of maternal cytoplasm, the residual body (RB) forms at the posterior end of dividing parasites and contains material remaining from the mother cell after daughter formation. Because no RB-specific molecular marker has yet been identified, the boundary between posterior maternal cytoplasm and a morphologically distinct nascent RB cannot be defined unambiguously at early stages of daughter formation. In this study, we therefore use the term “RB” to refer to the posterior compartment and connection between daughter parasites late during and after budding.

Increasing evidence indicates that the RB is not simply a terminal remnant of cell division but forms part of a dynamic system that maintains physical and functional continuity between parasites within the PV. An extensive F-actin network connects intravacuolar parasites through the RB and is required for parasite organization and material exchange (Periz et al., 2017a). Frénal et al. further demonstrated that intravacuolar tachyzoites are connected through their basal poles and the RB, allowing rapid exchange of soluble proteins between parasites and contributing to synchronized replication (Frenal et al., 2017). This functional connectivity depends on the unconventional myosins MyoI and MyoJ: deletion of either motor disrupts inter-parasite communication and replication synchrony (Frenal et al., 2017). Importantly, RBs are nevertheless still formed in the absence of MyoI or MyoJ, while the connection between the parasites and the RB is disrupted, demonstrating that RB formation can be uncoupled from functional connectivity (Frenal et al., 2017). This distinction is particularly important because it separates formation of the RB itself from its function as a conduit for inter-parasite exchange. The ability of the RB-associated network to mediate exchange between parasites raises the possibility that it may participate not only in the diffusion of soluble material but also in the redistribution of larger cellular components. Indeed, we previously demonstrated that maternal micronemal proteins, such as MIC2, are extensively recycled into daughter cells in an F-actin-dependent manner and can traffic through the intravacuolar network (IVN) (Periz et al., 2019). Together, these observations suggest that the cytoplasmic continuity provided by the RB and its associated F-actin network could contribute to the redistribution of maternal organelles during replication.

MyoF provides an additional link between myosin-dependent transport and organelle inheritance. MyoF is required for centrosome positioning and apicoplast inheritance, and its depletion causes enlargement of the RB accompanied by accumulation of micronemes and rhoptries (Jacot et al., 2013). Although this established a link between MyoF, RB morphology, and secretory-organelle distribution, whether the accumulated organelles represent maternally inherited material and whether they can subsequently be redistributed to daughter parasites remained unresolved. In addition, the actin-nucleating factors Formin-2 and Formin-3 contribute to organization of the intravacuolar F-actin network (Stortz et al., 2019; Tosetti et al., 2019). Together, these studies establish an actomyosin-dependent system connecting daughter parasites through the RB, but how this system contributes to the inheritance and redistribution of maternal organelles remains unclear.

F-actin could mediate this process either by serving as a track for myosin-dependent vesicular transport, as described for myosin VI in other eukaryotes (Frank et al., 2004), or through the intrinsic mobility and dynamic association of actin bundles themselves, which could transiently interact to promote vesicle exchange (Das et al., 2021; Khaitlina, 2014; Moore et al., 2021). In this study, we employ a dual-labeling strategy using the HaloTag system (Urh and Rosenberg, 2012) to investigate the fate of maternal organelles during T. gondii replication. Through fluorescence intensity-based analysis, we identify three distinct modes of organelle inheritance: inheritance of intact maternal organelles, expansion and division of pre-existing maternal organelles with incorporation of newly synthesized components, and degradation of maternal structures followed by de novo assembly. We further show that MyoF is required for efficient redistribution of maternal micronemes and rhoptries and that maternal secretory organelles accumulated in the RB after MyoF depletion can re-enter daughter parasites following restoration of MyoF. Together, these findings support a role for the RB as a dynamic compartment in the recycling and redistribution of maternal organelles.

## Results

### HaloTag-Based pulse chase labelling allows discrimination of recycled and de novo material

To evaluate HaloTag as a tool for distinguishing maternal from newly synthesized proteins during endodyogeny, two marker proteins with known fates were used: MIC2, which undergoes extensive recycling (Periz et al., 2019), and IMC1, primarily synthesized *de novo* (Ouologuem and Roos, 2014). Using CRISPR/Cas9, endogenous loci were tagged to generate MIC2-Halo and IMC1-Halo strains (Singer et al., 2023).

Both strains were sequentially labeled by incubation with membrane permeable dyes for 1h: maternal proteins were marked with Halo JF646 before invasion (M-MIC2, M-IMC1), and 24 hours later after replication, by Halo JF549 for 1h to label newly synthesized proteins (n-MIC2, n-IMC1) (Figure 1A). As *Toxoplasma gondii* replicates in an asynchronous manner, after 24 hours of replication, vacuoles containing 1, 2, 4, and 8 parasites can be observed in a single dish.

**Figure 1.**
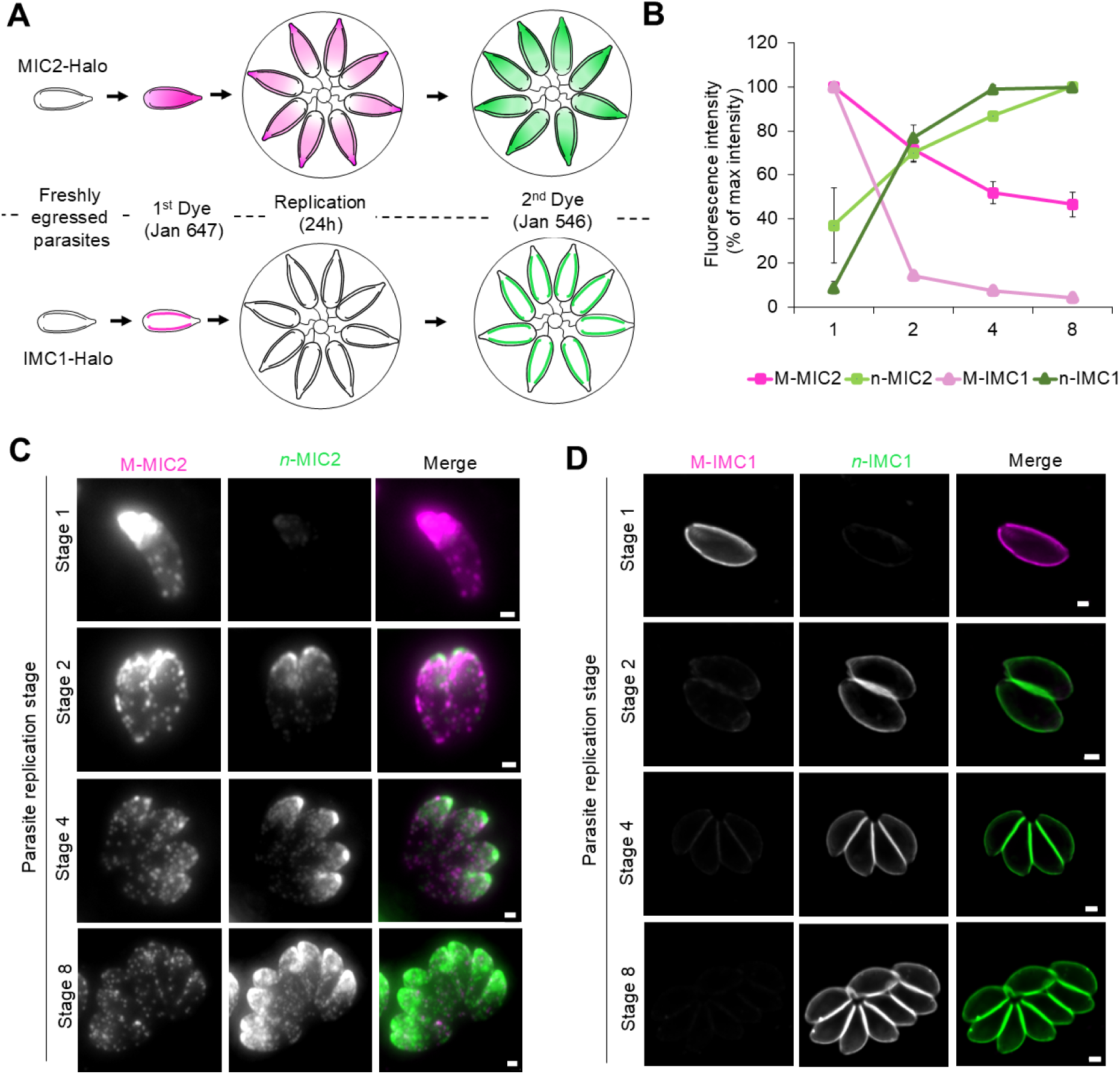
Discrimination between maternal and *de novo* material. **A)** Dual staining scheme. Maternal material is labelled initially with Janila Fluor®-646, and the excess washed out. After replication of parasites *de novo* synthesised material is labelled with Janila Fluor®-549, allowing efficient discrimination of the two populations. **B)** Fluorescence intensity quantification of maternal (M) and *de novo* sythenetised (n) MIC2 and IMC1 during replication from stage 1 to 8. Magenta: M-MIC2, Rosa: M-IMC1, Green: n-MIC2, Dark Green: n-IMC1. **C)** Representative picture of MIC2-Halo parasites during replication (stage 1 to 8) stained for both maternal (M) and *de-novo* (n) MIC2. Magenta: M-MIC2, Green: n-MIC2. Maternal MIC2 is efficiently recycled into the daughters, while de novo MIC2 is formed after each replication cycle **D)** Representative picture of IMC1-Halo parasites during replication (stage 1 to 8) stained for both maternal (M) and *de-novo* (n) IMC1. Magenta: M-IMC1, Green: n-IMC1. In contrast to micronemes, maternal IMC is degraded, and only residual amount is detected after the first replication cycle. Three biological replicates were used for all analysis with a total of 300 IMC-1 Halo vacuoles and 260 MIC2-Halo vacuoles analysed; error bars are standard deviations, and the centre measurement of the graph bars is the mean. Panels C and D show maximum-intensity projections of Z-stack images. All scale bars = 1 µm.

Fluorescence intensity (FI) analysis across those replication stages (Figure 1B–D, S1) revealed a progressive decline of M-MIC2 and an increase in n-MIC2, confirming protein recycling. In contrast, M-IMC1 FI dropped from 100% to 14% between stages 1 and 2, with a corresponding rise in n-IMC1 to ∼80%, indicating predominant de novo synthesis. These distinct FI dynamics demonstrate HaloTag’s utility for resolving protein origin and support the *de novo* assembly of the IMC.

### HaloTag profiling reveals three distinct patterns of protein inheritance

To further investigate protein inheritance during endodyogeny, endogenous HaloTag fusions were generated for a set of organelle-associated proteins: RON2 and ROP1 (rhoptries) (Besteiro et al., 2009; Striepen et al., 2001), GRA1 (dense granules) (Carruthers and Sibley, 1997), TIC20 (apicoplast) (van Dooren et al., 2008), SortLR (Golgi) (Sloves et al., 2012), ANKER1 (Barylyuk et al., 2020), GAPM1a (IMC) (Harding et al., 2019), and MyoA (glideosome) (Meissner et al., 2002) (Figure 2A, Figure S2).

**Figure 2.**
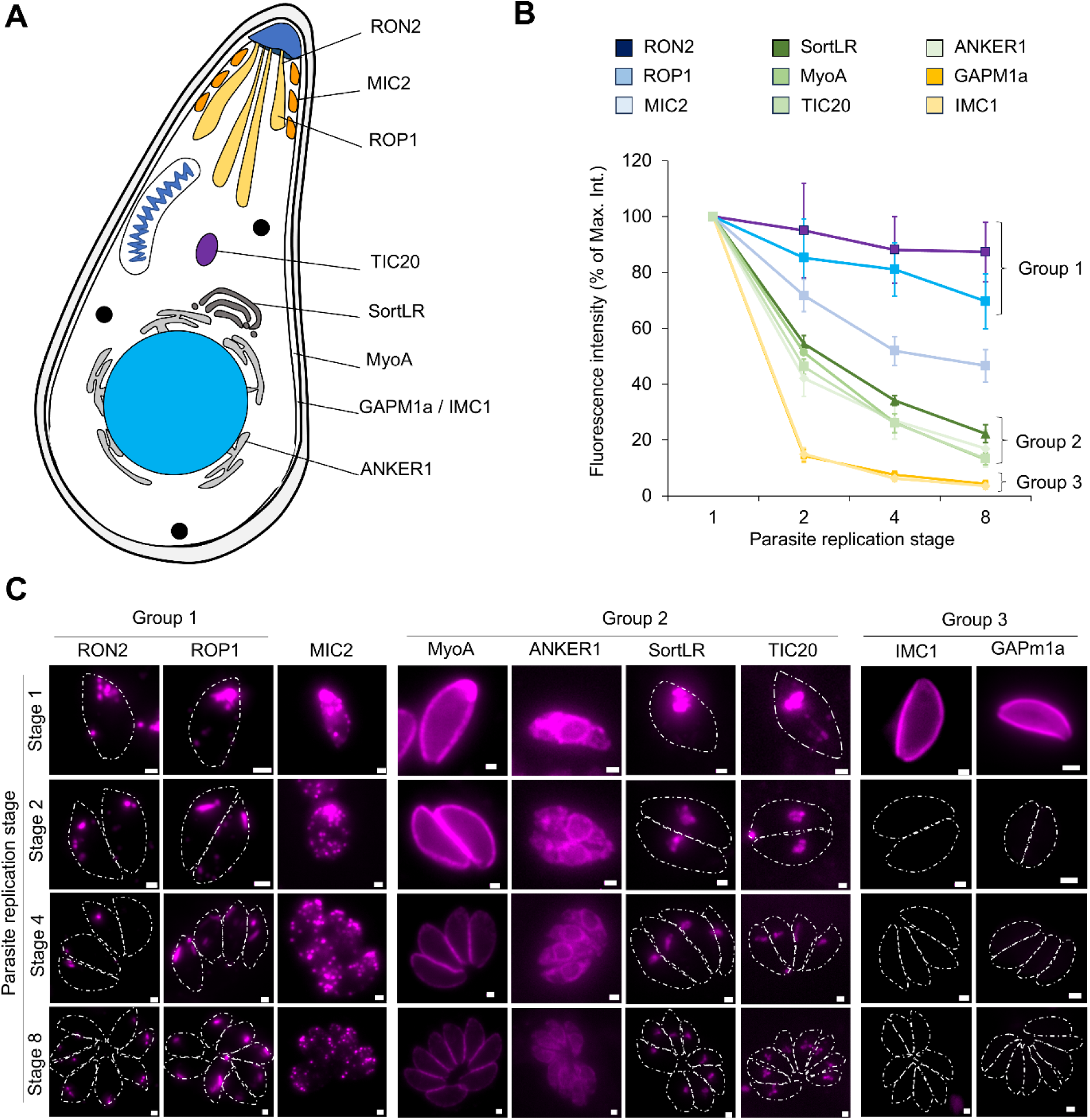
Three recycling fates for the maternal organelles. **A)** Overview of T. gondii organelles and molecular markers used for visualisation with endogenous Halo-Tags. **B)** Fluorescence intensity (FI) quantification of maternal (M) molecular marker listed in A. The fluorescence intensity variation during replication, allows to group them into three groups: 1) Efficient, almost quantitative recycling. 2) even distribution of maternal material and 3) almost exclusive *de novo* synthesis. Dark blue: RON2, Blue: ROP1, Pale blue: MIC2, Dark green: SortLR, Green: MyoA, Pale green: Tic20, Greenish white: ANKER1, Gold: GAPM1a, Yellow: IMC1. **C)** Representative picture of parasites expressing indicated Halo tagged proteins (stage 1 to 8) stained for maternal (M) proteins. Three biological replicates were used for all analysis with an average of 265 vacuoles analysed per protein; the graph indicate the mean value of FI calculated from the triplicate. Panels C shows maximum-intensity projections of Z-stack images. All scale bars = 1 µm.

Following the same sequential labeling strategy used for MIC2 and IMC1, fluorescence intensity (FI) of maternal proteins was quantified across the different replication stages (Figure 2B, C, S2C-G). Analysis revealed three distinct inheritance profiles:

- **Group 1:** RON2 and ROP1 exhibited stable maternal FI, with minimal decline of FI within a single rhoptry, indicating recycling of whole organelles. In good agreement with this hypothesis, detailed analysis of M-RON2 inheritance demonstrated that the maternal rhoptries are separated during the distribution and that some daughter cells did not obtain maternal organelles after successive rounds of replication (Figure 3D).
- **Group 2:** Proteins from diverse organelles (SortLR, ANKER1, TIC20, MyoA) showed a stepwise ∼50% reduction in maternal FI per division, suggesting that recycled, maternal material and *de novo* material end up in the same organelle
- **Group 3:** IMC proteins (IMC1 and GAPM1a) displayed a sharp FI loss between the first and second replication cycles, consistent with predominant *de novo* synthesis of the IMC and degradation of maternal material.

**Figure 3.**
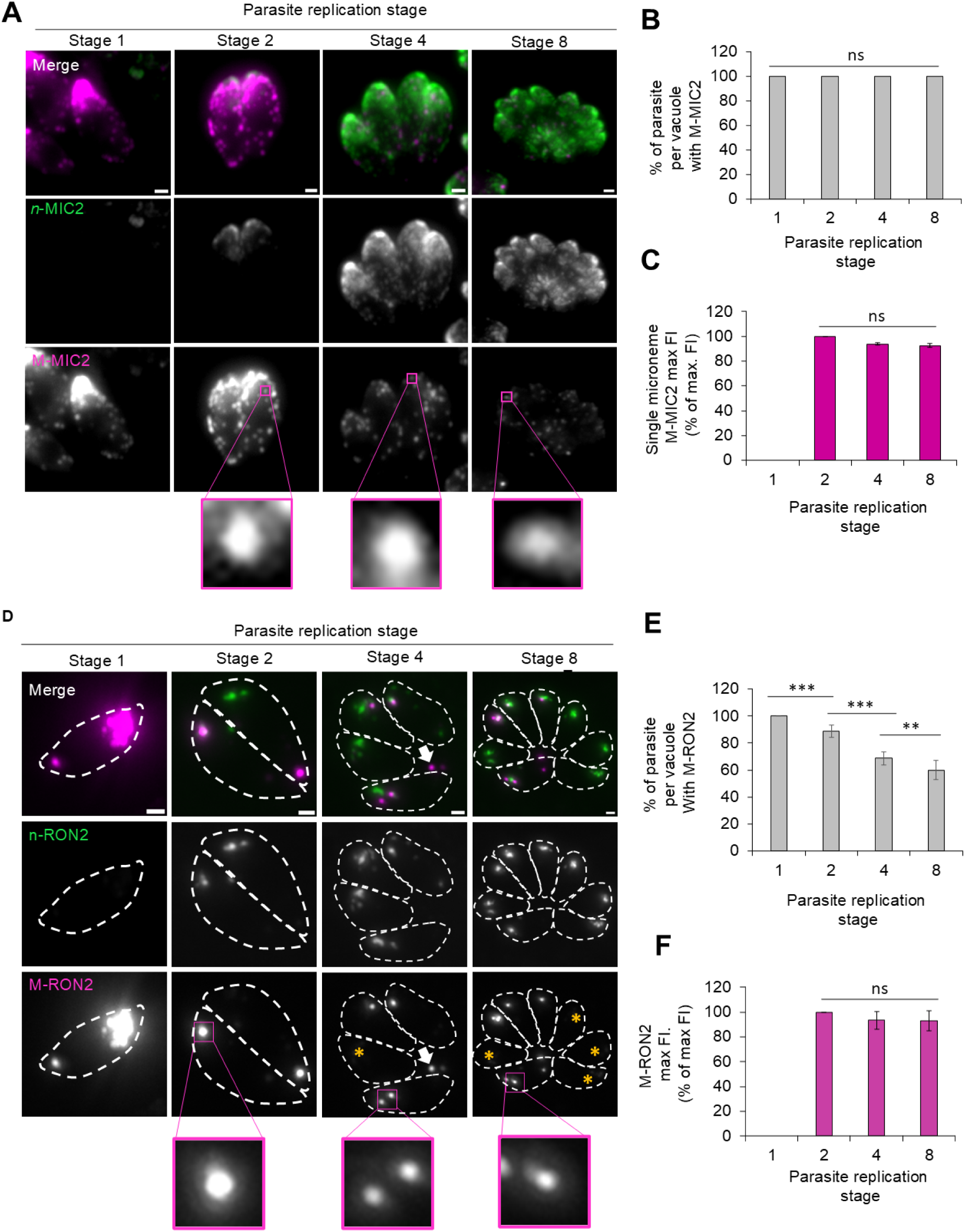
Micronemes and rhoptries are recycled as whole organelles. Representative picture of MIC2-Halo during replication (stage 1 to 8) stained for both the maternal (M) and *de novo* (n) MIC2. Magenta: M-MIC2, Green: n-MIC2. Zoom windows allow to visualise isolated apical M-MIC2 signal for which the fluorescence intensity is quantified in C. **B)** Quantification of the inheritance of M-MIC2 by the daughter cells during replication (stage 1 to 8). A total of 260 vacuoles were analysed. All daughter cells inherited M-MIC2. **C)** Fluorescence intensity quantification of isolated M-MIC2 signal as illustrated in A. A total of 203 isolated M-MIC2 signals were analysed. **D)** Representative picture of RON2-Halo during replication (stage 1 to 8) stained for both the maternal (M) and *de novo* (n) MIC2. Magenta: M-RON2, Green: n-RON2. Zoom windows allow to visualise isolated apical M-RON2 signal for which the fluorescence intensity is quantified in F. **E)** Quantification of the inheritance of M-RON2 by the daughter cells during replication (stage 1 to 8). A total of 289 vacuoles were analysed. Not all daughter cells inherit M-RON2 (asterisks), best seen in later replication stages. **F)** Fluorescence intensity quantification of isolated M-RON2 signal as illustrated in D. A total of 215 isolated M-RON2 signals were analysed. Three biological replicates were used for all analysis; error bars are standard deviations, and the centre measurement of the graph bars is the mean. All p-values ≤ 0.001 (***), using two-tailed unpaired Student’s t-test. Panels A and D show maximum-intensity projections of Z-stack images. All scale bars = 1 µm.

Interestingly, MIC2 exhibited intermediate behavior, prompting further analysis to refine its classification. These patterns highlight distinct organelle-specific protein inheritance mechanisms in *T. gondii*.

### Micronemes and rhoptries are recycled as intact organelles during Toxoplasma endodyogeny

Micronemes are known to comprise distinct subpopulations (Kremer et al., 2013), and prior work has shown that recycled and de novo MIC2 localize to separate microneme subsets (Periz et al., 2019). Based on these observations, we hypothesized that both micronemes and rhoptries are recycled as intact organelles, with new organelles formed independently via *de novo* synthesis.

To test this, we conducted detailed analyses of MIC2 and rhoptry inheritance. As expected, recycled (M-MIC2) and newly synthesized MIC2 (n-MIC2) were largely segregated into distinct micronemes with minimal colocalization (Figure 3A, S3)(Periz et al., 2019). Focusing on sparsely distributed micronemes outside the apical region (Figure 3A, boxed area), we measured fluorescence intensity (FI) of individual M-MIC2-positive micronemes at replication stages 2, 4, and 8 (Figure 3A, zoomed region; 3C). The relatively stable FI of these isolated structures supports whole organelle recycling, consistent with the FI profiles of rhoptries (Figure 2B, C).

Rhoptry analysis further corroborated this model. As with MIC2, M-RON2 and n-RON2 localized to distinct rhoptry populations (Figure S3), with divergence becoming more apparent at stage 8 (Figure 3D, Figure S4A). M-RON2 FI remained relatively constant through replication (Figure 3 F), while n-RON2 gradually increased from stage 2 onward (Figure S4B). Due to their low copy number (8–12 per tachyzoite), maternal rhoptries were separated and often absent in some daughter cells after successive divisions, consistent with stochastic whole-organelle inheritance illustrated by decrease in the average area of the M-RON2 fluorescence signal and the constancy of the total surface area (Figure S4C, D). As a consequence, a progressive decline in M-RON2-positive parasites per vacuole can be observed (Figure 3E, D; Figure S4A).

For the micronemes, even in larger parasitophorous vacuoles, all daughter parasites retained a mix of recycled and newly synthesized micronemes (Figure S5). At the 8-cell stage, maternal micronemes were distributed relatively evenly among daughter parasites (Figure S5A, B), and similar patterns were observed across independent vacuoles. This distribution is consistent with a non-random or regulated partitioning process, although the present data do not establish the underlying mechanism or imply the existence of a dedicated microneme-counting system. The simple probabilistic model for later-stage vacuoles further illustrates that highly even inheritance would be unlikely under unrestricted random segregation, but because the quantitative experimental dataset was obtained at the 8-cell stage, we interpret this analysis as supportive rather than definitive evidence for regulated partitioning. Together, these results support inheritance of micronemes and rhoptries as intact organelles, with *de novo* organellogenesis acting in parallel to replenish the secretory-organelle complement of daughter cells.

### Golgi inheritance involves coordinated expansion and partitioning of maternal and de novo proteins

Next, we analyzed SortLR, a Golgi marker, using dual HaloTag labeling. Maternal SortLR (M-SortLR) fluorescence declined by ∼50% per replication round, while newly synthesized SortLR (n-SortLR) increased proportionally, without abrupt loss (Figure 4A–B). This gradual dilution contrasts sharply with the rapid decay seen in Group 3 proteins.

**Figure 4.**
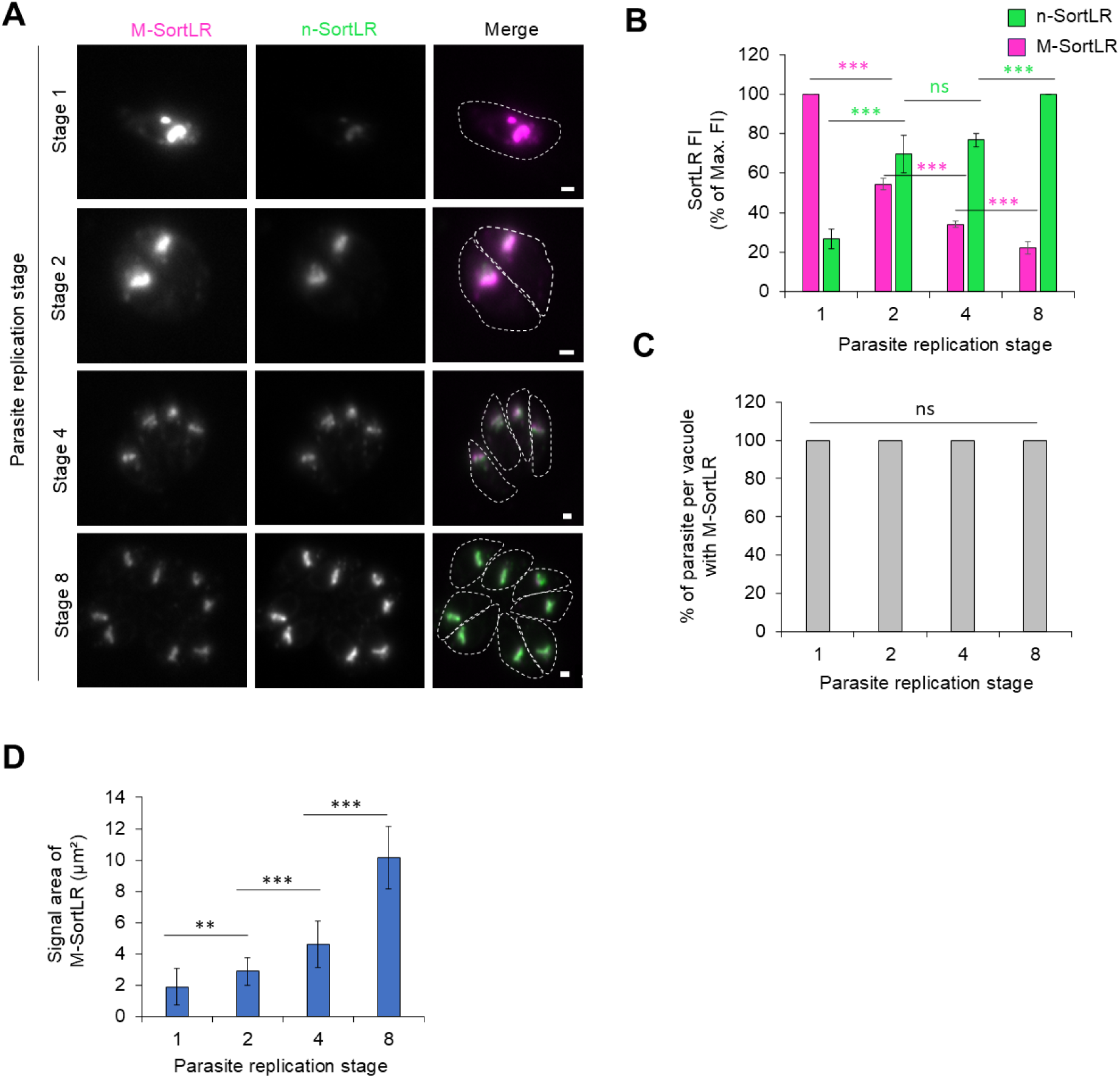
Expansion of the mother organelle for equivalent sharing to the daughter. **A)** Representative picture of SortLR-Halo during replication (stage 1 to 8) stained for both the maternal (M) and *de novo* (n) SortLR. Magenta: M-SortLR, Green: n-SortLR. **B)** Fluorescence intensity quantification of M-SortLR and n-SortLR illustrated in A. Magenta: M-SortLR, Green: n-SortLR. A total of 251 vacuoles were analysed. **C)** Quantification of the inheritance of M-SortLR by the daughter cells during replication (stage 1 to 8). All daughter cells inherited M-SortLR. A total of 100 vacuoles were analysed. **D)** Quantification of the signal area of M-SortLR during replication. A total of 100 vacuoles were analysed. The total surface of M-SortLR increase at each replication by about a factor 2. Three biological replicates were used for all analysis; error bars are standard deviations, and the centre measurement of the graph bars is the mean. All p-values ≤ 0.001 (***), using two-tailed unpaired Student’s t-test. Panels A shows maximum-intensity projections of Z-stack images. All scale bars = 1 µm.

M-SortLR and n-SortLR exhibited strong colocalization across replication stages, and all daughter cells retained M-SortLR (Figure 4C). The total Golgi area expanded from ∼2 µm² to ∼10 µm² by stage 8 (Figure 4D), consistent with coordinated Golgi elongation and medial fission, as described previously (Pelletier et al., 2002). These results demonstrate that the Golgi undergoes duplication, incorporating both maternal and *de novo* proteins during its growth and partitioning. Similar signal evolution was observed for the ER, MyoA and apicoplast (Figure S2, S6). Thus, Group 2 organelles follow a distinct inheritance mode, involving regulated expansion, integration of new material, and equal distribution to progeny—distinct from the whole-organelle recycling of rhoptries/micronemes and the complete turnover seen in IMC proteins.

### Maternal IMC proteins are degraded during replication

Group 3 proteins, typified by IMC components such as GAPM1a and IMC1, exhibit a sharply distinct inheritance profile, characterized by rapid loss of maternal fluorescence during replication. Using HaloTag-based dual labeling, we observed a ∼90% reduction in maternal GAPM1a signal by stage 2, with minimal retention through later stages (Figure 5A-C), indicative of degradation rather than recycling. M-GAPM1a and n-GAPM1a did not colocalize, and residual maternal signal localized transiently at the posterior pole before disappearing entirely (Figure 5D). This posterior accumulation occurs in the region that gives rise to the RB; however, because no marker uniquely distinguishes the nascent RB from the posterior maternal compartment, we cannot determine precisely when degradation occurs relative to RB formation.

**Figure 5.**
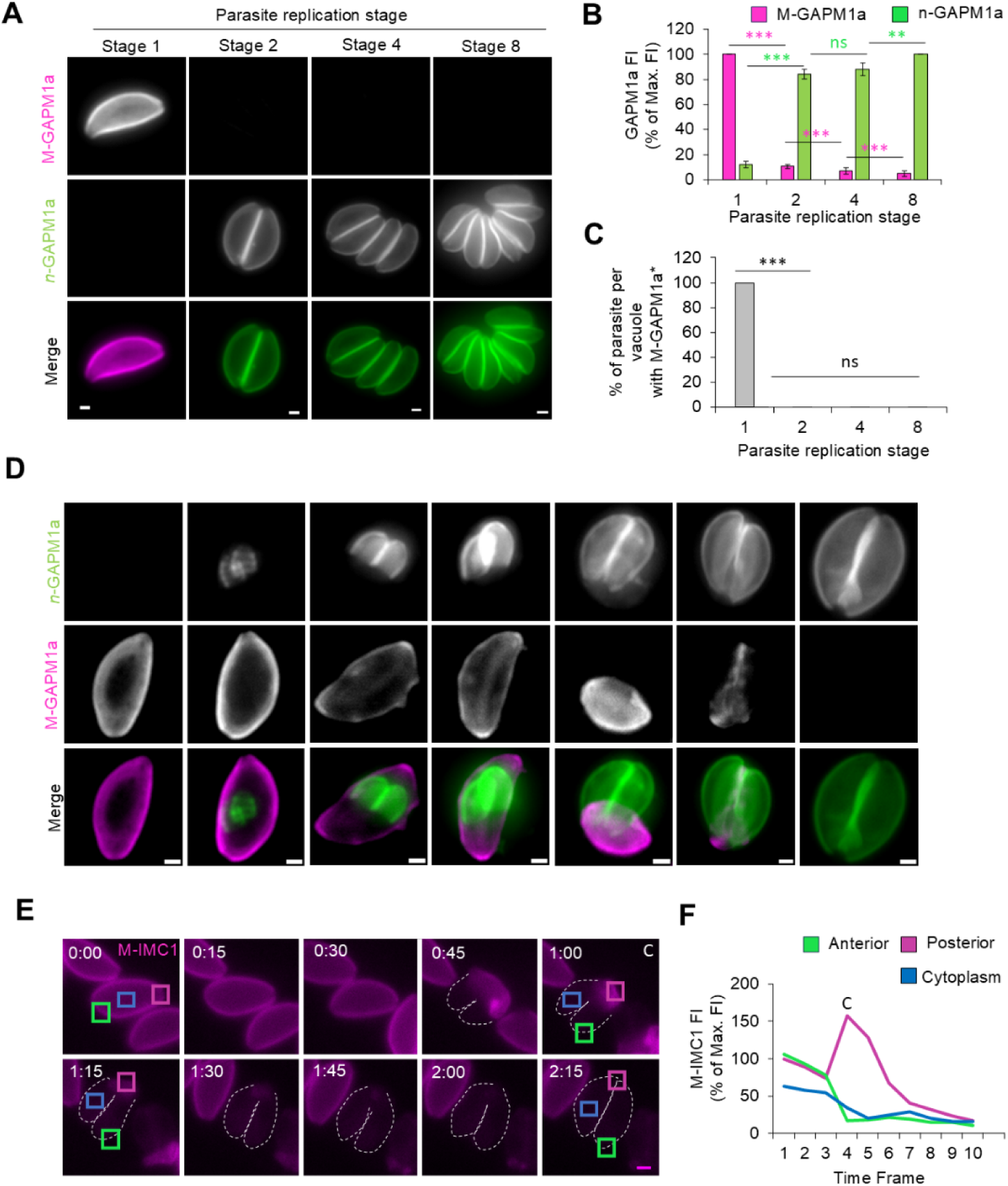
Degradation of the inner membrane complex. Representative picture of GAPM1a-Halo during replication (stage 1 to 8) stained for both the maternal (M) and *de novo* (n) GAPM1a. **B)** Fluorescence intensity quantification of M-GAPM1a and n-GAPM1a illustrated in A. A total of 265 vacuoles were analysed. **C)** Quantification of the inheritance of M-GAPM1a by the daughter cells during replication (stage 1 to 8). After replication, M-GAPM1a is not visible in daughter cells. A total of 265 vacuoles were analysed. *The calculation was performed using single stage parasite intensity to set up the exposure as performed for all the other organelles. Under those conditions the remaining signal of the maternal GAPM1a is below the detection level and cannot be observed. **D)** GAPM1a is degraded in the RB. Representative images of GAPM1a-Halo parasites at different stages of daughter cell development. M-GAPM1a collapses toward the forming residual body, where the signal disappears after completion of replication, indicating it’s degradation. **E)** Time series of IMC1-Halo parasites during the first replication. Parasite were stained for the maternal (M) IMC1 prior invasion. Three regions of the parasites were analysed: the apical (green ROI), the cytoplasmic (blue ROI) and the basal (magenta ROI). **F)** Fluorescence intensity analysis of the three regions defined in E. The curve and analysed ROI share the same colour code. Green: Apical, Blue: Cytoplasm, Magenta: Basal. After the accumulation of the mother IMC at the basal pole, the FI of M-IMC1 decrease without redistributing to any other region. Three biological replicates were used for all analysis; error bars are standard deviations, and the centre measurement of the graph bars is the mean. All p-values ≤ 0.001 (***), using two-tailed unpaired Student’s t-test. Panels A and D show maximum-intensity projections of Z-stack images. Panel E displays single optical plane images acquired during live imaging. All scale bars = 1 µm.

Live-cell imaging of IMC1 further supported this model: maternal IMC1 (M-IMC1) condensed at the posterior pole during daughter budding, followed by a marked fluorescence drop, without redistribution to other regions (Figure 5E, F and Figure S7, Video S1). This signal loss was replication-specific, as non-dividing parasites retained stable M-IMC1 levels (Figure S7D). Quantification across regions revealed a transient increase at the posterior, followed by complete disappearance, consistent with degradation (Figure 5F, Figure S7C).

Interestingly, DCs IMCs exhibited brighter fluorescence than the maternal IMC (Figure S7E, F), especially during early stages of elongation (Figure S7G, H). FI peaked when DCs reached ∼3 μm in length, suggesting the IMC is fully assembled before emergence and subsequently unfolds without requiring additional material. This preformation explains the absence of maternal IMC recycling. These findings align with earlier observations that IMC components are synthesized de novo during elongation (Ouologuem and Roos, 2014).

### Whole-organelle inheritance of micronemes and rhoptries occurs via similar recycling pathways

Our data reveal distinct spatial behaviors among the analysed organelle markers. Maternal microneme material can enter the posterior RB-associated compartment and subsequently redistribute to daughters, whereas most maternal RON2 is incorporated into daughter rhoptries before mother-cell collapse and formation of a morphologically distinct RB. In contrast, we did not observe comparable accumulation of the analysed Golgi, ER, or apicoplast markers in the RB during inheritance. Thus, the present experiments support participation of the RB in secretory-organelle redistribution but do not establish obligatory RB transit for all inherited micronemes or rhoptries.

To investigate this further, we performed live-cell imaging of rhoptry and microneme inheritance (Figure 6A, B, Video S2, S3). MIC2-Halo parasites co-expressing IMC1-YFP showed maternal MIC2 (M-MIC2) entering the posterior RB-associated compartment (Figure 6A, white arrows). After endodyogeny was completed, M-MIC2 present in this compartment was subsequently redistributed to the apical region of daughter cells (Figure 6A, 11:30, 16:00 h). These observations demonstrate that maternal micronemes can transit through the RB, which can act as a temporary reservoir from which they are subsequently redistributed to daughter parasites (Periz et al., 2019).

**Figure 6.**
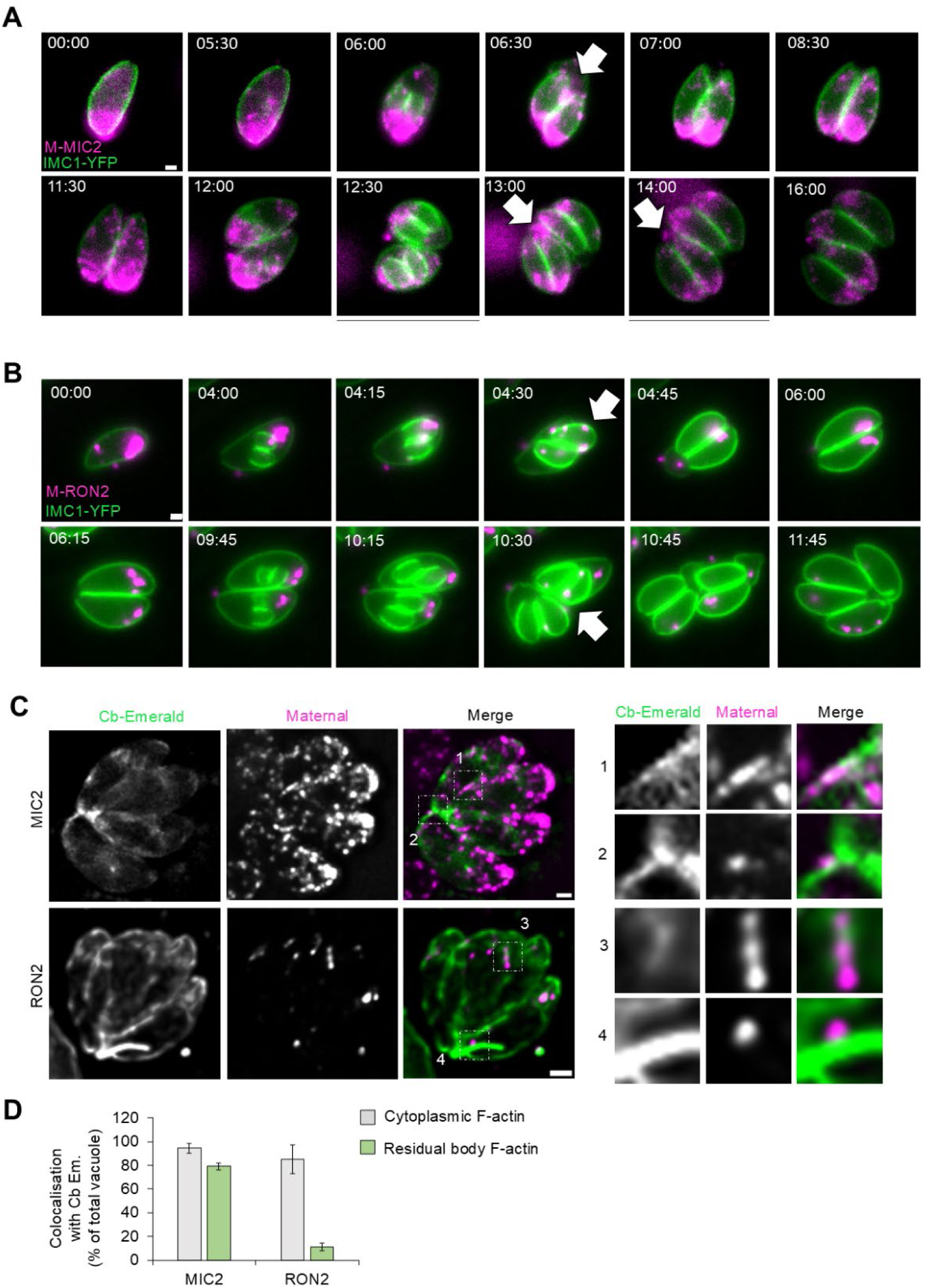
Micronemes and rhoptries are recycled similarly, but differ in timing and location. A) Time-lapse of MIC2-Halo over two replication cycles. Maternal MIC2 (M-MIC2) is stained, and daughter cell formation is visualized via IMC1-YFP. M-MIC2 is transported via the residual body (RB) (white arrows; see Video S2). B) Time-lapse of RON2-Halo parasites with IMC-YFP. Maternal RON2 (M-RON2) is transported before mother cell collapse and RB formation (white arrows). C) Representative images showing maternal organelles associating with F-actin and the RB. Insets highlight colocalization of maternal proteins and F-actin. D) Quantification of M-MIC2 and M-RON2 colocalization with F-actin. Data from three biological replicates, a total of 534 RON2-Halo and 342 MIC2-Halo vacuoles were analysed; bars represent means ± SD. Panel A and B display single optical plane images acquired during live imaging. Panel C show single Z plane of Z-stack images. Scale bars = 1 µm.

For rhoptries, we examined the inheritance of maternal RON2 (M-RON2) in RON2-Halo parasites co-expressing IMC1-YFP. In approximately 90% of parasites undergoing replication, M-RON2 was integrated into daughter rhoptries prior to mother-cell collapse and formation of a morphologically distinct RB (Figure 6B, 4:15-4:30 h and 10:30-10:45 h). M-RON2 was only occasionally detected in the RB-associated compartment. Thus, while both maternal micronemes and rhoptries can associate with the RB, their trafficking differs in timing, with most maternal RON2 being incorporated into daughter rhoptries before formation of a morphologically distinct RB. This observation also leaves open the possibility that a substantial fraction of maternal rhoptries is transferred directly from the mother into developing daughters. To test whether recycled rhoptries associate with F-actin, as previously reported for micronemes (Periz et al., 2019), we stably integrated the F-actin marker Cb-Emerald (Periz et al., 2017b) in the RON2-Halo line (Figure 6C, D). While m-MIC2 displayed clear F-actin– associated movement along filaments in the residual body (Figure 6C), colocalization of M-RON2 with Cb-Emerald in the residual body was infrequent. On the other hand, both secretory organelles were found to be associated with cytoplasmic F-actin (Figure 6D).

In summary, both rhoptries and micronemes can be inherited as intact organelles during *T. gondii* replication. Their distinct timing and localization indicate that related cytoskeletal trafficking mechanisms may operate through more than one spatial route during daughter-cell formation.

Given the F-actin dependency of microneme recycling demonstrated previously (Periz et al., 2019), we hypothesized that a conserved apicomplexan myosin contributes to this process. MyoF, a class XXII myosin conserved across Apicomplexa (Jacot et al., 2013), has been implicated in centrosome positioning, apicoplast inheritance, actin organization, and vesicular/endomembrane dynamics (Carmeille et al., 2021; Heaslip et al., 2016; Kellermeier and Heaslip, 2024). Interestingly, Jacot et al. (2013) previously reported enlargement of the residual body and accumulation of micronemes and rhoptries following MyoF depletion. The dual-labeling approach used here allows us to distinguish whether the accumulated material derives from the initial maternal protein pool or from proteins synthesized during subsequent replication cycles.

We endogenously Halo-tagged MIC2, RON2, and SortLR in the auxin-inducible degron line MyoF-mAID (Carmeille et al., 2021) and performed dual HaloTag labelling to differentiate maternal (M) from de novo (n) protein pools. Live imaging revealed that MyoF depletion resulted in accumulation of maternal MIC2 and RON2 in the RB-associated compartment during replication (Figure 7A, B). As Toxoplasma replicates asynchronously, different replication stages coexist within the same dish after 24 h. The second labeling step marks proteins synthesized since the beginning of the experiment, allowing discrimination between proteins present in the initial mother parasite and proteins synthesized during subsequent replication cycles. During successive rounds, proteins synthesized in earlier cycles can themselves become inherited material. Under MyoF depletion, these pools accumulate in the RB-associated compartment rather than being efficiently redistributed to daughters (Figure 7). Immunofluorescence with α-AMA1, α-MIC4, and α-MIC8 confirmed that this phenotype affects multiple micronemal proteins (Figures S8, S9). Approximately 90% of vacuoles exhibited RB-associated accumulation of maternal material upon auxin treatment (Figure 7D), demonstrating that MyoF is required for efficient redistribution of maternal micronemes and rhoptries. Interestingly, although our time-lapse analysis indicated that rhoptries only occasionally traffic through the RB, both maternal RON2 and MIC2 accumulate in the RB following MyoF depletion. This shared phenotype indicates that MyoF contributes to the redistribution of both organelle populations, even though their precise trafficking routes and timing may differ.

**Figure 7.**
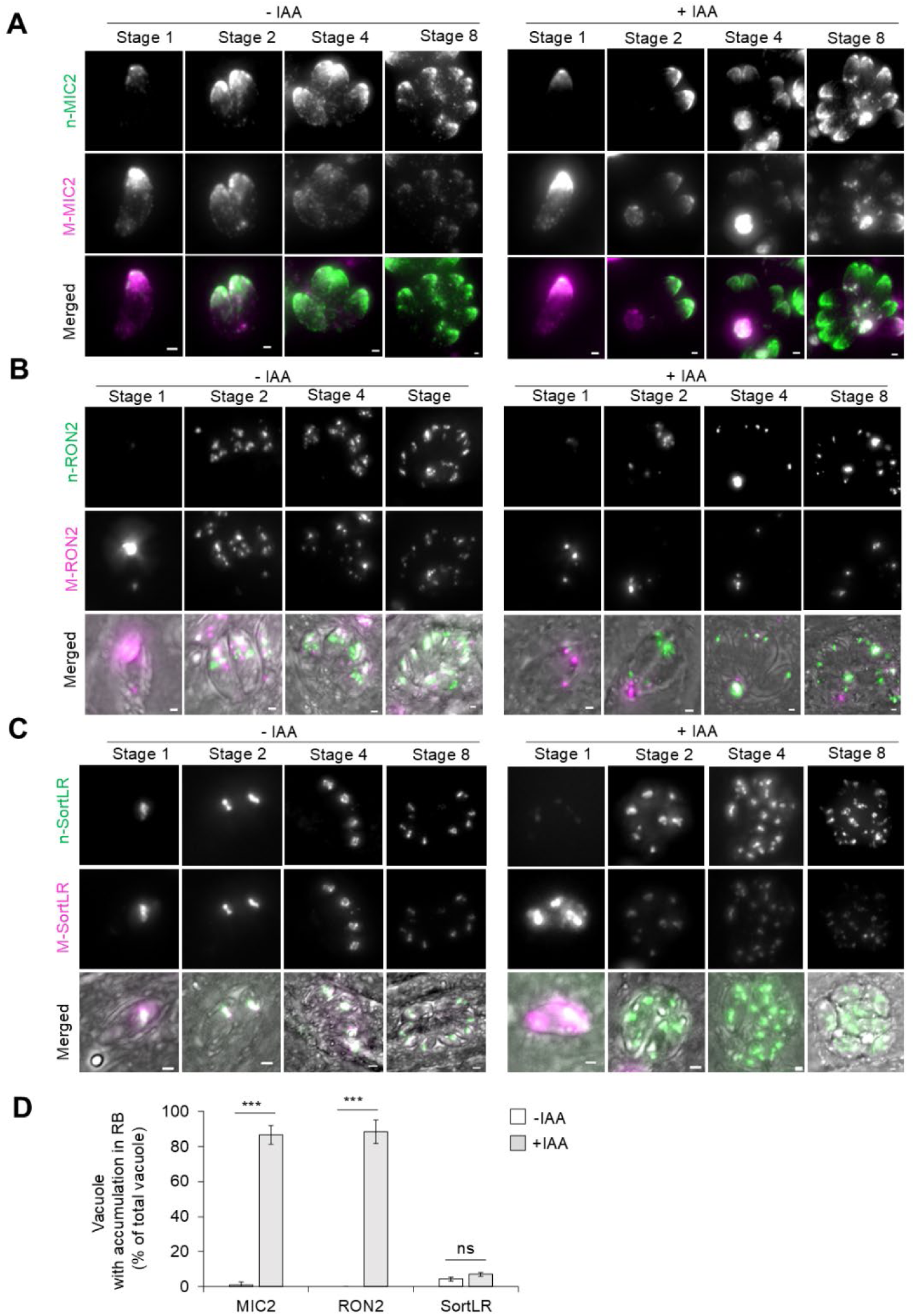
Myosin-F drives F-actin mediated recycling of maternal organelles via the residual body. A) Effect of MyoF knockdown (KD) on maternal microneme inheritance. MyoF-mAID MIC2-Halo parasites were labeled and grown ± auxin for 24h. Magenta: maternal MIC2 (M-MIC2), Green: newly synthesized MIC2 (n-MIC2). Without MyoF, M-MIC2 accumulates in the residual body (RB) instead of being passed to daughter cells (Stage 1–2), with increasing accumulation over replication cycles (Stage 2–8). B) Effect of MyoF KD on maternal rhoptry inheritance. MyoF-mAID RON2-Halo parasites show M-RON2 retention in the RB, mirroring the pattern seen with MIC2. C) Effect of MyoF KD on maternal Golgi inheritance. MyoF-mAID SortLR-Halo parasites show no significant RB accumulation of M-SortLR, unlike MIC2 and RON2. D) Quantification of vacuoles showing RB accumulation. White: control; Gray: MyoF-KD. Three biological replicates were used; a total of 675 Ctrl and 743 KD vacuoles were analyzed for MIC2-Halo, 645 Ctrl and 594 KD for RON2-Halo and 795 Ctrl and 619 KD for SortLR-Halo, bars show means ± SD. All p-values ≤ 0.001 (***), using two-tailed unpaired Student’s t-test. Panels A, B and C show maximum-intensity projections of Z-stack images. Scale bars = 1 µm

In contrast, Golgi inheritance (marked by SortLR) showed no comparable RB accumulation after MyoF depletion (Figure 7C). Neither accumulation in the RB nor separation of maternal and *de novo* SortLR signals was observed, consistent with previous reports that Golgi duplicates and partitions independently (Pelletier et al., 2002)), although Golgi fragmentation was observed as described previously.

To confirm that this phenotype results directly from MyoF depletion and not broader disruption of endomembrane architecture, we performed a rescue experiment. MIC2-Halo MyoF-mAID parasites were cultured in auxin for 24 hours to induce M-MIC2 retention in the RB-associated compartment, followed by auxin washout and continued replication (Figure 8A). Following auxin washout, M-MIC2 was redistributed to daughter parasites during subsequent replication cycles, although occasionally in an uneven manner (Figures 8A, 8C, and S10A). To determine whether this redistribution correlated with restoration of MyoF expression, we monitored MyoF recovery by IFA using the HA epitope fused to MyoF-mAID (Figure 8B). Following auxin washout, MyoF expression was restored. Quantification revealed a strong association between detectable MyoF and redistribution of maternal micronemes (Figure S10B), supporting a requirement for MyoF in efficient retrieval/redistribution of accumulated maternal MIC2.

**Figure 8.**
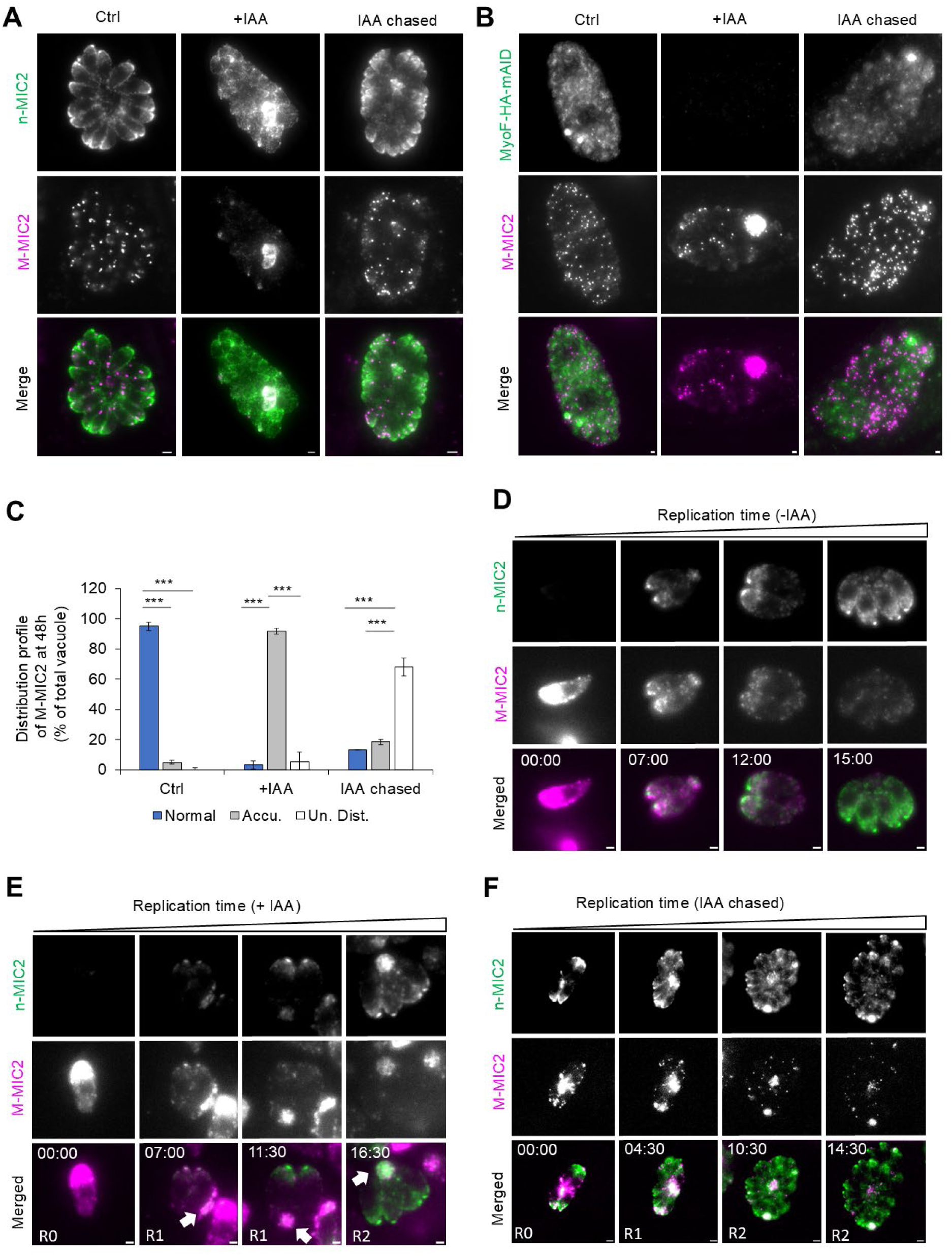
The residual body (RB) functions as a recycling center, not a dead end. To assess the fate of material accumulated in the RB, auxin chase experiments were performed: 24h with auxin followed by 24h without. A) Representative images of MyoF-mAID MIC2-Halo parasites after 48h in control (no auxin), continuous auxin (Aux), or auxin washout (Aux washed). Magenta: M-MIC2, Green: n-MIC2. Continuous auxin led to M-MIC2 accumulation in the RB, while auxin washout enabled redistribution. B) Representative images of MyoF-mAID MIC2-Halo parasites after 48h in control (no auxin), continuous auxin (Aux), or auxin washout (Aux washed). Magenta: M-MIC2, Green: α-HA (MyoF). In absence of auxin MyoF is visible, induction with auxin deplete MyoF led to M-MIC2 accumulation in the RB, while auxin washout enabled MyoF production and M-MIC2 redistribution. C) Quantification of vacuoles showing normal, accumulated, or redistributed M-MIC2. White: control; Gray: MyoF-KD; Blue: MyoF-KD auxin chase. D) Time-lapse without auxin: M-MIC2 briefly passes through the RB (white arrows). A total of 170 Ctrl, 186 KD and 179 auxin chased vacuoles were analyzed. E) Time-lapse with auxin: sustained M-MIC2 accumulation in the RB throughout replication. F) Time-lapse after auxin washout: initial M-MIC2 accumulation in the RB resolves by the second replication cycle, with redistribution observed. Three biological replicates were used; bars show means ± SD. All p-values ≤ 0.001 (***), using two-tailed unpaired Student’s t-test. Panels A and B show maximum-intensity projections of Z-stack images. Panel D, E and F display single optical plane images acquired during live imaging. Scale bars = 1 µm.

Live-cell imaging confirmed these dynamics (Videos S4-S6). In untreated parasites, M-MIC2 entered the RB-associated compartment and was later redistributed to daughter cells (Figure 8D, Video S4). Under auxin treatment, M-MIC2 accumulated post-budding and failed to redistribute efficiently (Figure 8E; Video S5). Following auxin washout, imaging was initiated after removal of auxin at 24 h; time 0 in Figure 8F/Video S6 therefore corresponds to the start of post-washout imaging. Redistribution was observed during the subsequent replication cycle rather than immediately after MyoF re-expression (Figure 8F; Video S6). This delay suggests that efficient redistribution requires sufficient recovery of MyoF function and/or progression into the next replication cycle.

To assess whether microneme degradation occurred during retention within the residual body, the fluorescence intensity of individual maternal micronemes was compared between control vacuoles and vacuoles subjected to MyoF depletion followed by auxin washout (Figure S10C, D). No significant difference in microneme fluorescence intensity was detected between control micronemes and micronemes that had remained within the RB for 24 h prior to redistribution. These results indicate that micronemes recovered after retention within the RB do not show a detectable loss of maternal MIC2 signal. Together, these findings establish that MyoF is required for the efficient recycling and redistribution of maternal micronemes and rhoptries. They further demonstrate that the RB can function as a dynamic compartment from which retained maternal organelles can be redistributed to daughter parasites.

## Discussion

### Use of dual labelling to differentiate maternal and de novo material

Understanding the complexity of organelle inheritance in *Toxoplasma gondii* requires tools capable of resolving both the spatial and temporal dynamics of protein trafficking. Conventional single-labelling approaches have been limited to static snapshots, obscuring key processes such as organelle biogenesis, recycling, and selective degradation. To overcome these challenges, we employed a dual HaloTag pulse-chase labelling strategy to distinguish maternally inherited from de novo–synthesized proteins. This approach, previously underutilized in apicomplexan parasites (Koreny et al., 2023; Periz et al., 2019), allowed us to resolve protein fates across replication cycles and map distinct inheritance routes for multiple organelles (Figure 9A).

**Figure 9.**
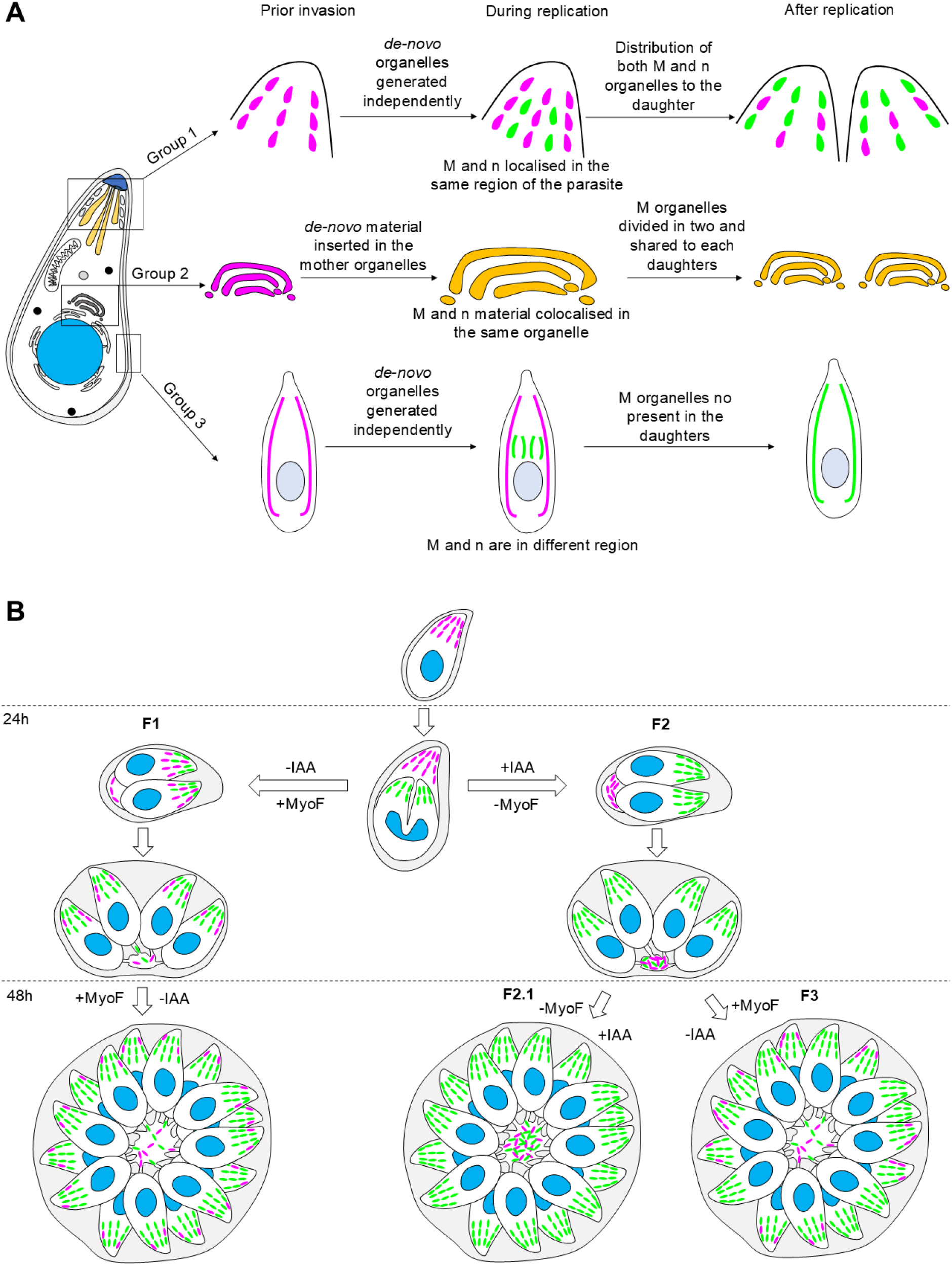
Schematic summaries. A: Summary of the three fates of the maternal organelles. Group 1: Efficient, intact organelle recycling. De novo organelles are generated independently of the maternal organelle but in a similar location. Both de novo and maternal are distributed to the daughter cells. The Fluorescence intensity of the maternal organelle is relatively stable. Group 2: Even distribution of maternal material through organelle expansion with the insertion of the de novo material in the mother organelle. The Fluorescence intensity of the maternal organelle is divided by two at each replication step but their signal surface is increased by two. Group 3: Almost exclusive de novo synthesis. The de novo organelle is generated independently of the maternal organelle and in a different location. The maternal organelle is not clearly observable after a cycle of replication. The fluorescence intensity of the maternal organelle drastically drops after the first replication. Magenta: maternal organelle, Green: de novo organelle, Yellow: Colocalization between maternal and de novo. B: Summary of the MyoF regulated inheritance. The maternal and the de novo micronemes are generated independently. In presence of MyoF (F1), the daughter cells form a chimera possessing both maternal and *de novo* micronemes, which are distributed in a relatively equal manner even after multiple replication cycles. In absence of MyoF, (F2) the inherited micronemes accumulate in the residual body and the daugthers cells are mostly composed of de novo micronemes. If the depletion of MyoF is maintained (F2.1) more and more inherited micronemes accumulate inside the residual body without important degradation. In opposition, if the expression of the MyoF is reestablished by the auxin removal (F3), the inherited micronemes stuck inside the residual body are redistributed but in a less equal manner than usually observed for control.

#### The residual body as a dynamic compartment for organelle recycling

Our findings reaffirm the emerging view of the RB as an active, multifunctional structure rather than a passive cytoplasmic remnant (Frenal et al., 2017; Periz et al., 2017a). First described over 50 years ago (Sheffield and Melton, 1968), the RB was long believed to serve as a repository for discarded material, often called a “waste bin” (Nishi et al., 2008). **Our** results extend this view by showing that maternal secretory organelles can enter the RB and subsequently be redistributed to daughter parasites in a MyoF dependent manner (Figure 9B). This is particularly evident for micronemes: live-cell imaging demonstrated that maternal MIC2 can transit through the RB, while MyoF depletion causes maternal micronemes to accumulate within this compartment. Restoration of MyoF allows this accumulated material to re-enter daughter parasites (Figure 8), demonstrating that the RB is not simply a terminal or degradative compartment but can serve as a temporary reservoir during organelle recycling. Rhoptry inheritance reveals a more complex picture. Although maternal RON2 was occasionally detected in the RB and accumulated prominently in this compartment following MyoF depletion, most maternal RON2 was incorporated into daughter rhoptries before mother-cell collapse and formation of a morphologically distinct RB. Thus, our data support a role for the RB in secretory-organelle redistribution but do not imply that transit through the RB is obligatory for all inherited micronemes or rhoptries. Direct transfer from the maternal parasite into developing daughters may operate in parallel, particularly for rhoptries. Importantly, the shared accumulation of maternal MIC2 and RON2 in the RB following MyoF depletion indicates that MyoF contributes to the redistribution of both organelle populations, even though their precise spatial routes and timing may differ.

This active role is further supported by the dependence of RB-mediated recycling on F-actin and the class XXII myosin MyoF, which we show to be essential for retrieval of maternal MIC2 and RON2 but dispensable for Golgi inheritance although we noticed a fragmentation of the Golgi, which has been described to depend on MyoF (Carmeille et al., 2021). The RB is also critical for cytoplasmic continuity across parasites, synchronizing replication and facilitating inter-parasite protein exchange (Frenal et al., 2017; Muniz-Hernandez et al., 2011; Periz et al., 2017a). Importantly, RB formation and functional connectivity can be uncoupled: residual bodies are still formed in the absence of MyoI or MyoJ, whereas parasites lose their connection to the RB and inter-parasite communication is disrupted (Frenal et al., 2017). This distinction suggests that formation of the RB alone is insufficient for its exchange functions and that actomyosin-dependent connectivity is required to establish a functional route between parasites.

Together with the F-actin network extending through the RB (Periz et al., 2017a), these observations support a model in which the RB forms part of a regulated system for exchange and redistribution of cellular material within the vacuole rather than simply representing a passive leftover of daughter-cell formation. Given these parallels, the RB may be functionally analogous to the mammalian midbody remnant, which regulates post-mitotic signaling and intracellular trafficking(Kuriyama et al., 2025; Peterman and Prekeris, 2019).

### Fate determination: inheritance or degradation?

An important question raised by our study is how the parasite determines whether maternal proteins are recycled or degraded. Our results indicate that this decision is selective rather than stochastic (Figure 9). For instance, micronemes recovered after retention in the RB following MyoF depletion retain maternal MIC2 fluorescence intensities comparable to control micronemes (Figure S10C, D). This indicates that the recovered organelles do not show detectable loss of maternal MIC2 signal. However, because this analysis measures individual recovered micronemes, it cannot exclude degradation of a fraction of the total microneme population during RB retention.

In contrast, IMC proteins like GAPM1a and IMC1 are lost within a single cycle and show no evidence of recycling. Both proteins transiently accumulate at the posterior pole before their maternal signal disappears raising questions about the underlying mechanism.

Although T. gondii encodes homologs of lysosomal and autophagic degradation machinery (Besteiro et al., 2011; Smith et al., 2021; Thaprawat et al., 2025), none have been localized to the RB or directly linked to IMC degradation. However, ubiquitin ligases and proteasomal components have been identified in the RB proteome (O’Shaughnessy et al., 2023; Que et al., 2002). IMC proteins are present in the ubiquitinated proteome (Silmon de Monerri et al., 2015), and recent work showed that deletion of the kinase ERK7 prevents sequestration of the E3 ligase CSAR1, resulting in aberrant degradation of conoids (O’Shaughnessy et al., 2023). These findings raise the possibility that selective ubiquitination and regulated proteolysis contribute to determining the fate of maternal proteins during daughter-cell formation. Whether these processes occur directly within the RB or within the posterior maternal compartment that subsequently forms the RB remains to be determined.

### A call to reassess recycling factors in Toxoplasma gondii

The recognition of the residual body (RB) as a key site for organelle recycling has fundamentally revised our understanding of intracellular trafficking in *T. gondii* (Periz et al., 2019; Tosetti et al., 2019). Earlier models emphasized *de novo* synthesis of organelles and proposed a repurposing of classical endocytic machinery for secretory functions (Tomavo, 2014). However, our HaloTag-based pulse-chase studies show that secretory organelles such as micronemes and rhoptries are extensively recycled during daughter-cell formation, with the RB serving as an important site for their redistribution in a MyoF-dependent manner.

This shift in our understanding calls for a systematic re-evaluation of trafficking factors traditionally implicated in protein targeting, vesicle transport, and organelle biogenesis. Many of these, particularly Rab-GTPases, SNAREs, and dynamins were previously studied without consideration of recycling pathways. Notably, accumulation of microneme material in the RB has been observed upon perturbation of several trafficking regulators, including Rab5A (Kremer et al., 2013), ArlX3 (Klinger et al., 2024), SORTLR (Sloves et al., 2012), VPS8 (Morlon-Guyot et al., 2018), and VPS11 (Morlon-Guyot et al., 2015). Yet, whether these phenotypes reflect a block in secretion, synthesis, or recycling remains unclear. To resolve this, a targeted reanalysis of trafficking factors using our dual-labelling HaloTag assay combined with the splitCas9 system (Li et al., 2022) would allow high-throughput functional screening to specifically distinguish defects in recycling from those in de novo biogenesis, offering a path to redefine the roles of classical and lineage-specific trafficking regulators in *T. gondii*.

## Material and Methods

### Parasite culture and genetic manipulation

#### Growth and generation of transgenic T. gondii

*T. gondii* tachyzoites from the RH strain and derived lines, including RH Δku80/TATi and RH Δku80/Tir1, were maintained at 37 °C with 5% CO₂ in human foreskin fibroblasts (HFFs; ATCC, SCRC 1041) cultured in Dulbecco’s Modified Eagle Medium (DMEM; Sigma, D6546) supplemented with 10% fetal bovine serum (FBS; BioSell, FBS.US.0500), 4 mM L-glutamate (Sigma, G7513), and 20 μg/mL gentamycin (Sigma, G1397) as previously described (Gras et al., 2019).

#### Generation of transgenic parasites

New strains were generated using CRISPR/Cas9 as previously described (Li et al., 2022). Guide RNAs (gRNAs) targeting the regions of interest were designed using EuPaGDT (Alvarez-Jarreta et al., 2024). All gRNA and primer sequences are listed in Supplementary Table 1. Briefly, gRNA oligos were annealed, ligated into the Cas9-YFP vector, and verified by sequencing (Eurofins Genomics). Repair templates were generated by PCR amplification of the Halo or YFP tag flanked by 50 bp homology arms to the target gene, using Q5 High-Fidelity DNA Polymerase (New England BioLabs). PCR products were purified with a PCR purification kit (Blirt, EM26.1). Tachyzoites were mixed with the repair template and 10 µg of the corresponding Cas9 vector, and transfected using the Amaxa 4D-Nucleofector system (Lonza, AAF-1003X). Transfected parasites were allowed to invade fresh HFFs and replicate for 48 h. Following manual egress and filtration through a 3 μm filter, Cas9-YFP-expressing parasites were enriched by FACS (FACSAria III, BD Biosciences) and sorted into 96-well plates. Correct integration of the repair template was confirmed by PCR.

### Labelling and characterization

#### Maternal and de-novo protein discrimination

Fresh tachyzoites expressing Halo-tagged reporters were mechanically egressed, filtered through a 3 μm filter, and resuspended in cold DMEM containing a membrane-permeable Halo dye, Janelia Fluor® 646 (1:1000, Promega), for 1 h. Parasites were centrifuged at 2,500 rpm for 5 min and washed three times with fresh medium to remove unbound dye. Parasites were then seeded onto HFF-covered Ibidi live-cell dishes for overnight replication. After replication, a second labeling was performed with a new membrane-permeable Halo dye, Janelia Fluor® 549 (1:1000, Promega) for 1 h at 37 °C, followed by three washes before imaging.

#### Fluorescence intensity across the replication

Parasites were labeled as described above and allowed to replicate for 24 h on HFF-coated Ibidi live-cell dishes. Approximately 15 fields of view were imaged using Z-stacks spanning 3 μm centered on the vacuoles (Figure S11A). For each replication stage (1, 2, 4, and 8 parasites per vacuole), individual tachyzoites were sampled across multiple vacuoles.

Maximum-intensity projections were generated from non-deconvolved images. Fluorescence intensity (FI) was quantified as the maximum gray value measured within regions of interest (ROIs) drawn on individual tachyzoites from each vacuole stages (Figure S11B). ROIs excluded overlapping parasites, neighboring vacuoles, and regions with atypical signal intensity (Figure S11C). For each biological replicate, up to 25 tachyzoites per replication stage were analyzed, and the mean FI value was calculated for each stage. The highest mean FI observed among the stages within a replicate was defined as 100%, and FI values for the other stages were expressed relative to this maximum. Relative FI values were then averaged across three independent biological replicates. Data are presented as mean ± SD. A total of 272, 286, 251, 276, 223, 284, 265, 300 and 260 vacuoles were analysed for RON1, ROP1, SortLR, MyoA, TIC20, ANKER1, GAPM1a, IMC1 and MIC2 respectively.

#### Fluorescence intensity of the daughter cells vs new mother cells

Images were processed as described above. Fluorescence intensity was quantified as the maximum gray value measured within regions of interest (ROIs) drawn on individual daughter cells and on newly formed mother cells. To reliably discriminate newly formed mother IMC from daughter IMC (both in the same channel colour, Jan. 549), the channel corresponding to the initial maternal IMC (Jan. 646) was intentionally overexposed, allowing unambiguous identification of the new mother cell IMC signal. ROIs were drawn on non-overlapping regions and excluded areas containing overlapping parasites, neighboring vacuoles, or atypical signal intensity, following the same exclusion criteria used for replication-stage measurements (Figure S11D). Quantification was performed on maximum-intensity projections generated from non-deconvolved Z-stacks. Data are presented as mean ± SD from vacuoles pooled across three independent biological replicates.

#### Measurement of the signal area of the fluorescence

To quantify the average and total fluorescent area, images were analyzed in Fiji. Up to 25 isolated vacuoles per stage (1 to 8) were cropped and analyzed individually. A threshold was applied to the maternal signal, and both average and total signal area were measured. Three independent biological replicates were performed, and values are reported as mean ± SD.

#### Quantification of percentage of parasite per vacuole with maternal protein

Images were analyzed in Fiji to determine the proportion of parasites per vacuole with maternal protein signal. Up to 25 vacuoles per stage (1 to 8) were cropped and analyzed individually. Vacuoles where all parasites retained maternal signal were scored as 100%. If some parasites lacked signal, the percentage of maternal-positive parasites was calculated. The experiment was performed in three independent biological replicates. Values are reported as mean ± SD.

#### Live replication assay

Parasites were labeled with Janelia Fluor® 646 (1:1000, Promega) for 1 h and washed three times before transfer to Ibidi live-cell dishes covered with HFFs. Parasites were allowed to invade for 1–2 h to obtain ∼10 parasites per field of view. Excess parasites were removed by washing three times. Imaging was performed on a Leica DMI8 microscope at 37 °C in 5% CO₂. Images were acquired every 15–30 min for 12 h using minimal laser power and exposure. Experiments were performed in three biological replicates and analyzed in Fiji.

#### Colocalization assays: ANKER1-Halo with HDEL-GFP

Parasites expressing ANKER1-Halo were transiently transfected with an HDEL-GFP plasmid to label the ER. After overnight replication in HFF-coated Ibidi dishes, parasites were labeled with Janelia Fluor® 646 (1:1000, Promega) for 1 h and washed three times prior to imaging. Z-stacks of HDEL-GFP-positive parasites were acquired. The proportion of vacuoles displaying ER localization of ANKER1-Halo was assessed manually in Fiji/ImageJ by inspection of full Z-stacks and maximum-intensity projections. In addition, the degree of colocalization between ANKER1-Halo and HDEL-GFP was quantified using Pearson’s correlation coefficient calculated with the Coloc2 plugin in Fiji/ImageJ. Pearson analysis was performed on 30 individual parasites obtained from three independent biological replicates (10 parasites per replicate). The mean Pearson’s correlation coefficient was 0.92 ± 0.04 (mean ± SD).

#### Colocalization assays: MIC2/RON2 with Cb-Emerald

For analysis of association with the F-actin network, MIC2 or RON2 was endogenously tagged and labeled in a strain stably expressing Cb-Emerald. Freshly egressed parasites were labeled with Janelia Fluor® 646 (1:1000, Promega) for 1 h, washed, and seeded onto HFF-coated live-cell dishes. After 24 h of replication, parasites were fixed in 4% paraformaldehyde and imaged. Z-stacks of vacuoles displaying a clearly defined F-actin network were acquired, and the association of MIC2- or RON2-positive structures with F-actin was assessed in both the parasite cytoplasm and the RB. Analysis was performed manually in Fiji/ImageJ by inspection of full Z-stacks and maximum-intensity projections. The RB was defined by the presence of thick bundles of F-actin located at the basal pole of the parasites and forming a continuous structure connecting parasites within the vacuole, consistent with the characteristic morphology of the RB. Cytoplasmic F-actin was defined as all actin signal located within the parasite body excluding this RB-associated region. Three independent biological replicates were analyzed, and data are reported as mean ± SD.

#### Quantification of the maternal microneme distribution

To assess the distribution of maternal micronemes at stage 8, parasites were labeled as described in “Maternal and de novo protein discrimination.” Twenty-five stage 8 vacuoles were selected. Using the de novo MIC2 signal, the outline of each tachyzoite was traced, and the number of maternal MIC2-positive parasites was counted. Tachyzoites were numbered 1 to 8 from left to right. Three biological replicates were performed. Data are reported as mean ± SD. Images were analyzed in Fiji.

#### Probability of maternal microneme inheritance

To explore whether the relatively even distribution of maternal micronemes could arise through stochastic segregation alone, we previously considered a simplified model allocating 64 micronemes among 32 daughter tachyzoites. Because the experimental quantification presented in Figure S5 was performed at the 8-cell stage, this 32-cell calculation is not used here as direct evidence for regulated segregation. A rigorous test of regulated partitioning will require a null model matched to the experimentally quantified stage and, ideally, direct quantification of later-stage vacuoles.

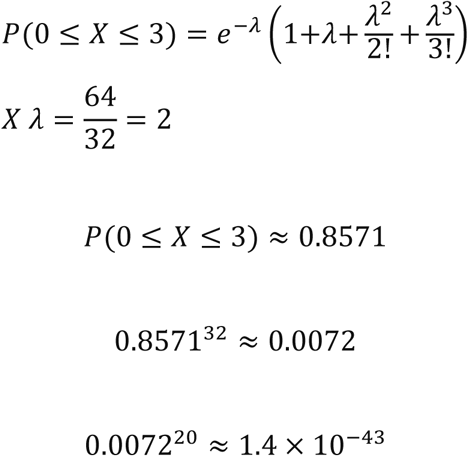

The relatively even maternal MIC2 distribution observed at the 8-cell stage is therefore described as an empirical observation rather than evidence for an established microneme-counting or balancing mechanism.

### Phenotypic assays

#### Induction of MyoF KD

MyoF-mAID parasites were genetically modified to endogenously tag MIC2, RON2 and SortLR. As previously described for MyoF mAID parasites (Carmeille et al., 2021), MyoF-mAID MIC2/RON2 or SortLR-Halo parasites were induced +/- auxin for 4h prior to labelling and experiments.

#### Quantification of the accumulation of protein in the residual body

Parasites were treated ± auxin for 4 h and labeled as described above. After 24 h replication on live-cell dishes, ∼100 vacuoles were imaged across ∼15 fields of view. The percentage of vacuoles showing accumulation of material in the residual body was calculated. The experiment was performed with three independent biological triplicates. The values were then expressed as the mean values of the three independent experiments ± SD. Images were analysed via Fiji.

#### Auxin chased experiment

Parasites were induced ± auxin for 4 h prior to labeling as described above in “Maternal and de novo protein discrimination”. Following transfer to live-cell dishes, parasites were allowed to replicate for 48 h in the continued presence or absence of auxin before imaging. For the auxin chase condition, parasites were first maintained in auxin-containing medium for 24 h. The medium was then removed, and dishes were washed three times with fresh medium to ensure complete auxin removal before addition of fresh medium lacking auxin. Parasites were subsequently allowed to replicate for an additional 24 h prior to imaging.

To assess recovery of MyoF expression following auxin washout, parallel samples were fixed in 4% paraformaldehyde and processed for immunofluorescence using an anti-HA antibody to detect the MyoF-mAID-HA fusion protein. Vacuoles were categorized according to the presence or absence of detectable MyoF signal and correlated with the corresponding microneme distribution phenotype.

Approximately 15 fields of view were imaged per condition, corresponding to ∼100 vacuoles. The percentage of vacuoles displaying microneme accumulation within the residual body was determined. In addition to the “accumulated” and “normal” phenotypes, a third category termed “uneven distribution” was defined for vacuoles in which micronemes were no longer retained within the residual body but had not yet reached the homogeneous distribution observed in control parasites. The frequency of each phenotype was quantified. Experiments were performed in three independent biological replicates, and values are presented as mean ± SD. Image analysis was performed using Fiji/ImageJ.

#### MyoF-mAID Live replication assay

Parasites were induced +/- auxin for 4 h prior to labelling. Parasites were labelled with the membrane-permeable Halo dye Janelia Fluor® 646 (1:1000, Promega) for 1 h and washed three times prior to transfer onto HFF-covered live-cell dishes. Parasites were allowed to invade for 1–2 h, after which excess parasites were removed by three washes. Janelia Fluor® 549 (1:10000, Promega) was added to the media to record de novo generation of the protein of interest during live imaging. Live-cell dishes were then transferred to the Leica DMI8 at 37 °C and 5% CO₂. Laser power and exposure were adjusted to the lowest values allowing reliable imaging. Images were taken every 15–30 min for 12 h to follow the tagged protein during replication. For the auxin-washout assay, auxin was removed after 24 h of replication as described above (“Auxin chased experiment”), and live imaging was subsequently initiated. Time 0 in Figure 8F and Video S6 corresponds to the first acquired frame after auxin washout. Experiments were performed in three independent biological replicates and images were analysed in Fiji.

### Imaging

#### Widefield microscopy

Unless stated otherwise, all images were acquired on a Leica-DMI8, objective 100x with the LasX software (v3.7.4). Fiji (v1.53c) was used to analyse the picture and all counts were made manually. LasX software (v.) from Leica was used to obtain parasite imaging data and All images and movies were processed using Fiji (ImageJ) software v Image Processing Software (Schindelin et al., 2012).

#### Software

Fiji (FIJI ImageJ v1.54f) was used to analyze the picture and all counts were made manually.

#### Data analysis

All data were plotted using Microsoft Excel

#### Statistical analysis

Two-tailed unpaired Student’s t-tests were performed to evaluate statistical significance.

- For Figures 2C, 2E, 2F, 3B, 3D, 4B, 4C, 5C (stage 1–2), and Supplementary Figures S1A, S1B, S2C–S2G, S4B–S4D, differences between stages 1–2, 2–4, and 4–8 were assessed.
- For Figures 2B, 4C, and 5C (stage 2–8), variance could not be calculated because values were constant (0 or 100%), and these were therefore labeled as non-significant.
- For Figures 7D, S8D, and S9B, differences between uninduced controls and MyoF knockdown (KD) induced samples were analyzed.
- For Figure 8C, the distribution of phenotypes (normal, accumulated, and redistributed micronemes) was compared between control, auxin-induced (48 h), and auxin-washed (chased) conditions.
- For Figure S5B, microneme distribution across positions 1–2, 2–3, 3–4, 4–5, 5–6, 6–7, and 7– 8 was analyzed; no statistically significant differences were observed, and all were labeled as non-significant.
- Finally, for Figure S7F, the difference in fluorescence intensity between daughter and mother IMC signals was assessed.

Data are presented as mean ± SD. Statistical significance ns: not significant (p ≥ 0.05); * : p < 0.05; ** : p < 0.01; *** : p < 0.001.

## Supporting information

Summary Data

## Acknowledgements

This work was supported by the Deutsche Forschungsgemeinschaft (DFG, German Research Foundation) grant GR 5696/2-1 to Simon Gras, grant ME 2675/6-2 and DFG Equipment grant INST 86/1831-1 to Markus Meissner. We thank VEuPathDB for their invaluable Informatics Resources. The MyoF-mAID strain was kindly provided by Prof. Aoife Heaslip (University of Conneticut), MIC4, MIC8 and AMA1 antibodies were kindly provided by Prof Domique Soldati-Favre (University of Geneva) and Prof. Gary Ward (University of Vermont) respectively. We thank Dr. Elena Jimenez-Ruiz for the useful discussions.

## Supplementary Figures

**Figure S1.**
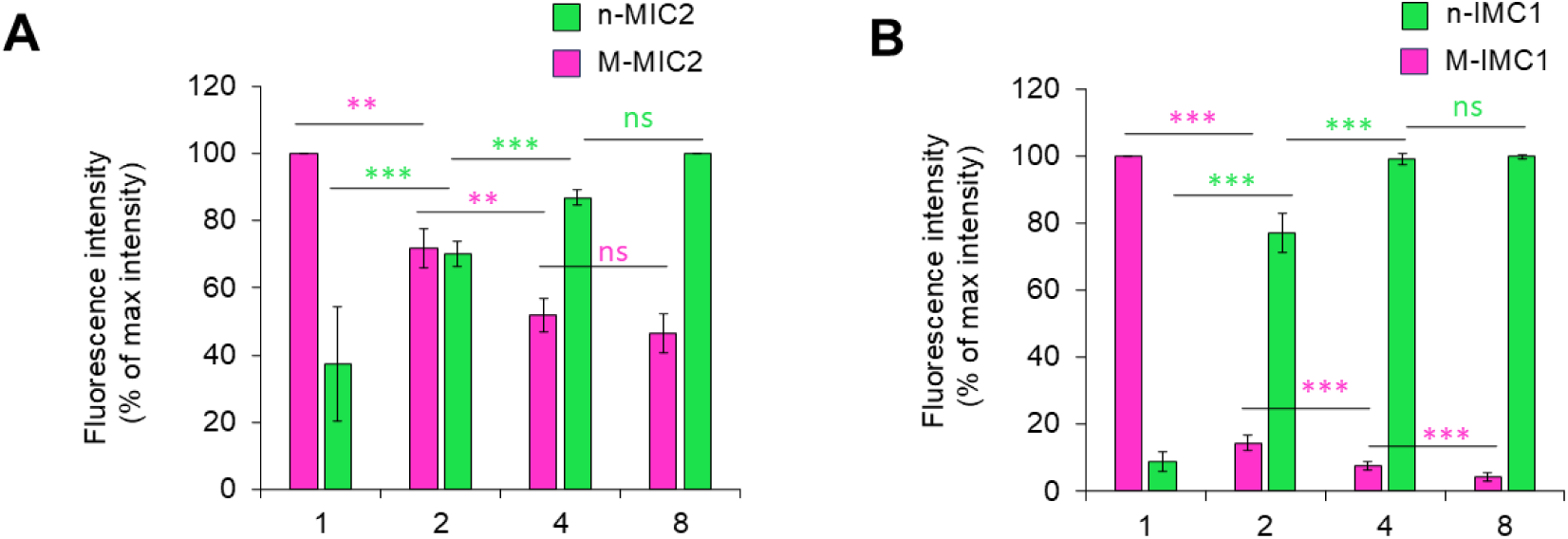
Fluorescence intensity of maternal and *de novo* MIC2 and IMC1. **A)** Fluorescence intensity quantification of M-MIC2 and n-MIC2 illustrated in Figure 1A, B and presented here independently and with statistical analysis. Magenta: M-MIC2, Green: n-MIC2. **B)** Fluorescence intensity quantification of M-IMC1 and n-IMC1 illustrated in Figure 1A, B and presented here independently and with statistical analysis. Magenta: M-IMC1, Green: n-IMC1. Three biological replicates were used for all analysis; a total of 300 and 260 vacuoles were analysed for IMC1 and MIC2 respectively, bars show means ± SD. All p-values ≤ 0.001 (***), using two-tailed unpaired Student’s t-test.

**Figure S2.**
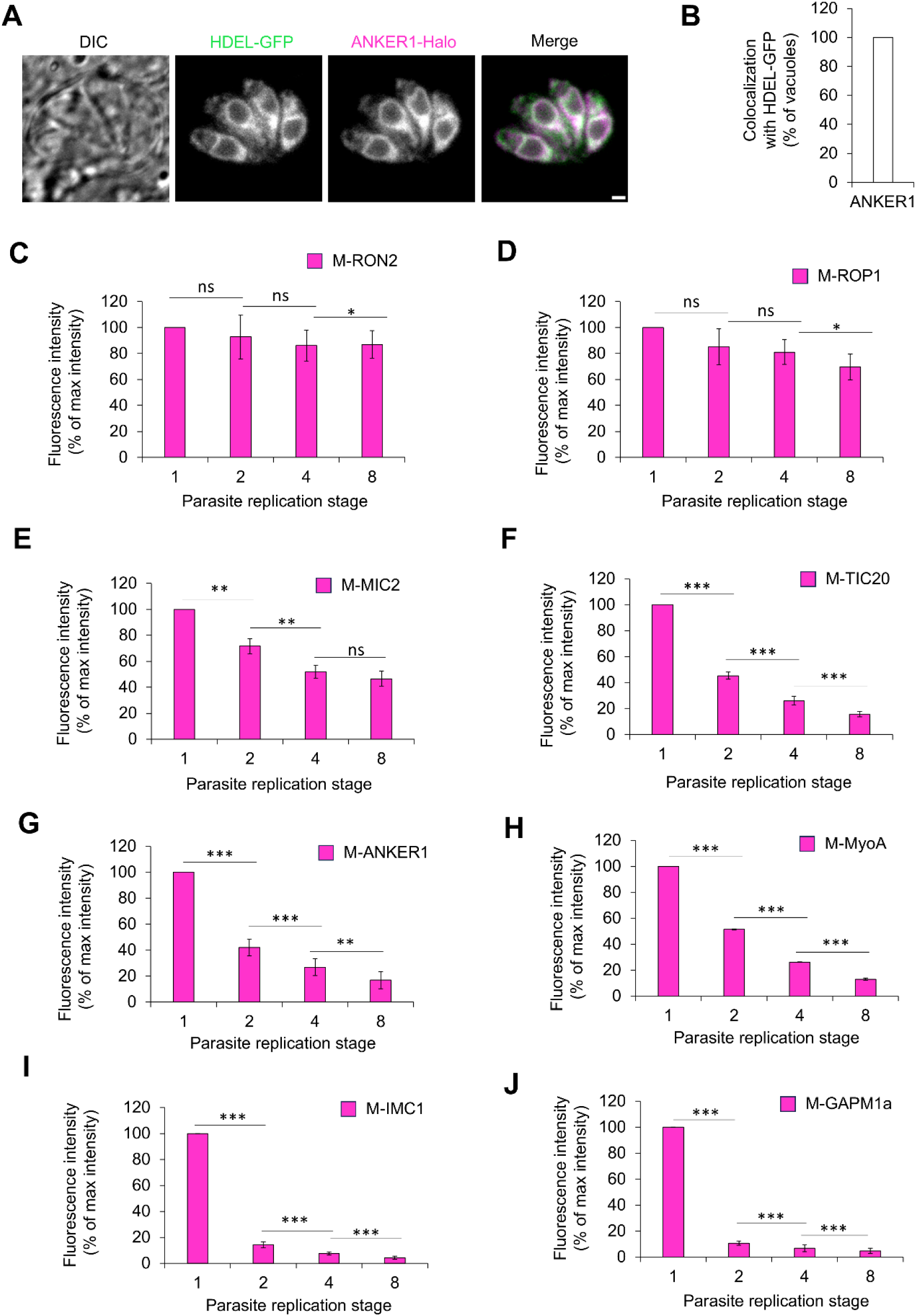
ANKER1 is a resident of the ER and independent fluorescence analysis from 1B. **A)** Representative picture of the colocalization ANKER1-Halo with HDEL-GFP transfected parasites. Magenta: ANKER1-Halo, Green: HDEL-GFP. **B)** Quantification of the percentage of vacuoles with colocalization between ANKER1-Halo and HDEL-GFP. A total of 169 vacuoles were analysed. Pearson correlation coefficient was also calculated between ANKER-1 and HDEL-GFP from a total 30 independent vacuoles, 0,92±0,02. **C)** Fluorescence intensity quantification of M-RON2 illustrated in Figure 2B and presented here independently and with statistical analysis. A total of 272 vacuoles were analysed **D)** Fluorescence intensity quantification of M-ROP1 illustrated in Figure 2B and presented here independently and with statistical analysis. A total of 286 vacuoles were analysed. **E)** Fluorescence intensity quantification of M-MIC2 illustrated in Figure 1B, 2B and S1A and presented here independently and with statistical analysis. A total of 260 vacuoles were analysed. **F)** Fluorescence intensity quantification of M-TIC20 illustrated in Figure 1B and presented here independently and with statistical analysis. A total of 223 vacuoles were analysed. **G)** Fluorescence intensity quantification of M-ANKER1 illustrated in Figure 1 B and presented here independently and with statistical analysis. A total of 284 vacuoles were analysed. **H)** Fluorescence intensity quantification of M-MyoA illustrated in Figure 1B and presented here independently and with statistical analysis. A total of 276 vacuoles were analysed. **I**) Fluorescence intensity quantification of M-IMC1 illustrated in Figure 1B, 2B and S1B and presented here independently and with statistical analysis. A total of 300 vacuoles were analysed. **J**) Fluorescence intensity quantification of M-GAPM1a illustrated in Figure 2B and 5B and presented here independently and with statistical analysis. A total of 265 vacuoles were analysed. All datasets were regrouped here to allow a better visual comparison between conditions. Three biological replicates were used for all analysis; error bars are standard deviations, and the centre measurement of the graph bars is the mean. All p-values ≤ 0.001 (***), using two-tailed unpaired Student’s t-test. Panels A shows maximum-intensity projections of Z-stack images. All scale bars = 1 µm.

**Figure S3.**
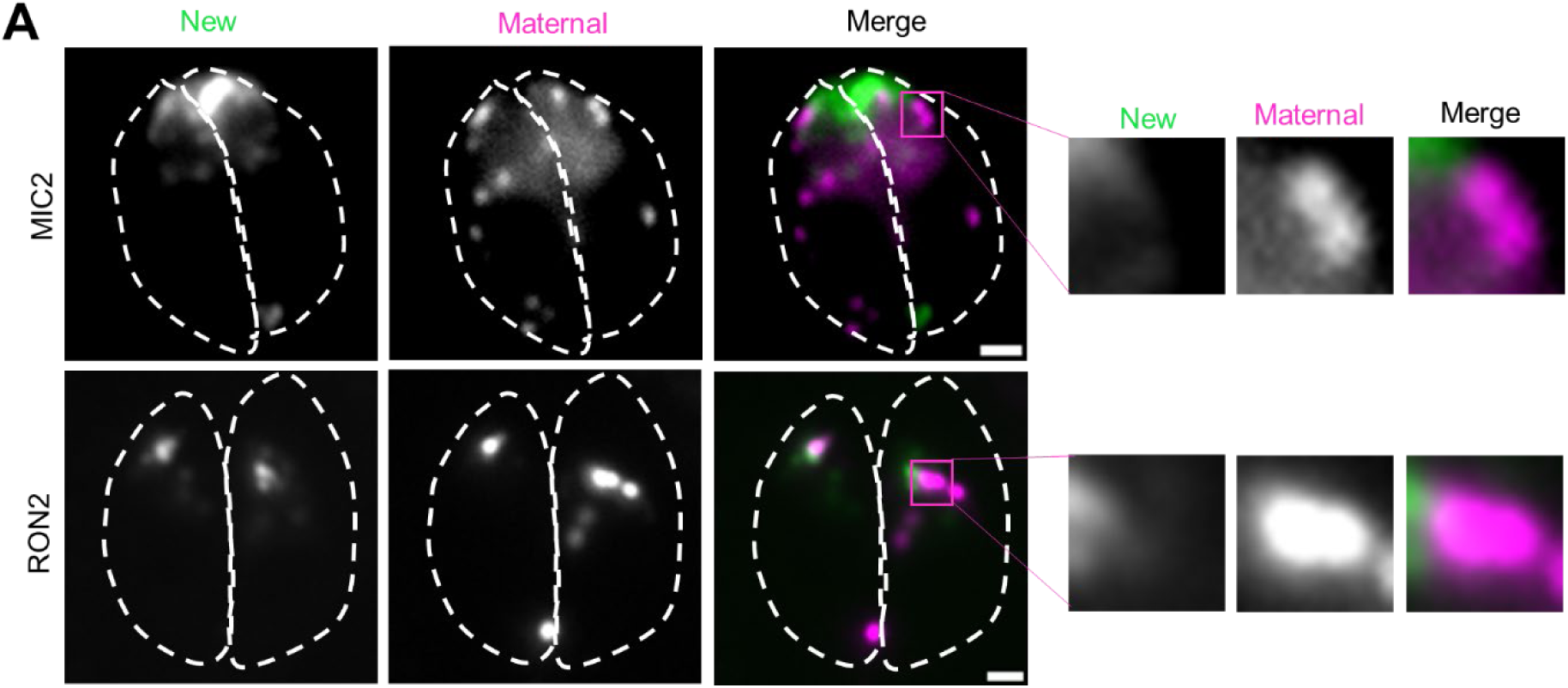
Rhoptries are inherited intact. **A)** The *de-novo* secretary organelle are generated independently of the maternal organelles. Representative picture of MIC2 and RON2-Halo parasites. Magenta: Maternal, Green: *de novo*. Zoom window highlight the absence of colocalization between maternal and de novo material. Three biological replicates were used. Panels A shows maximum-intensity projections of Z-stack images. All scale bars = 1 µm.

**Figure S4.**
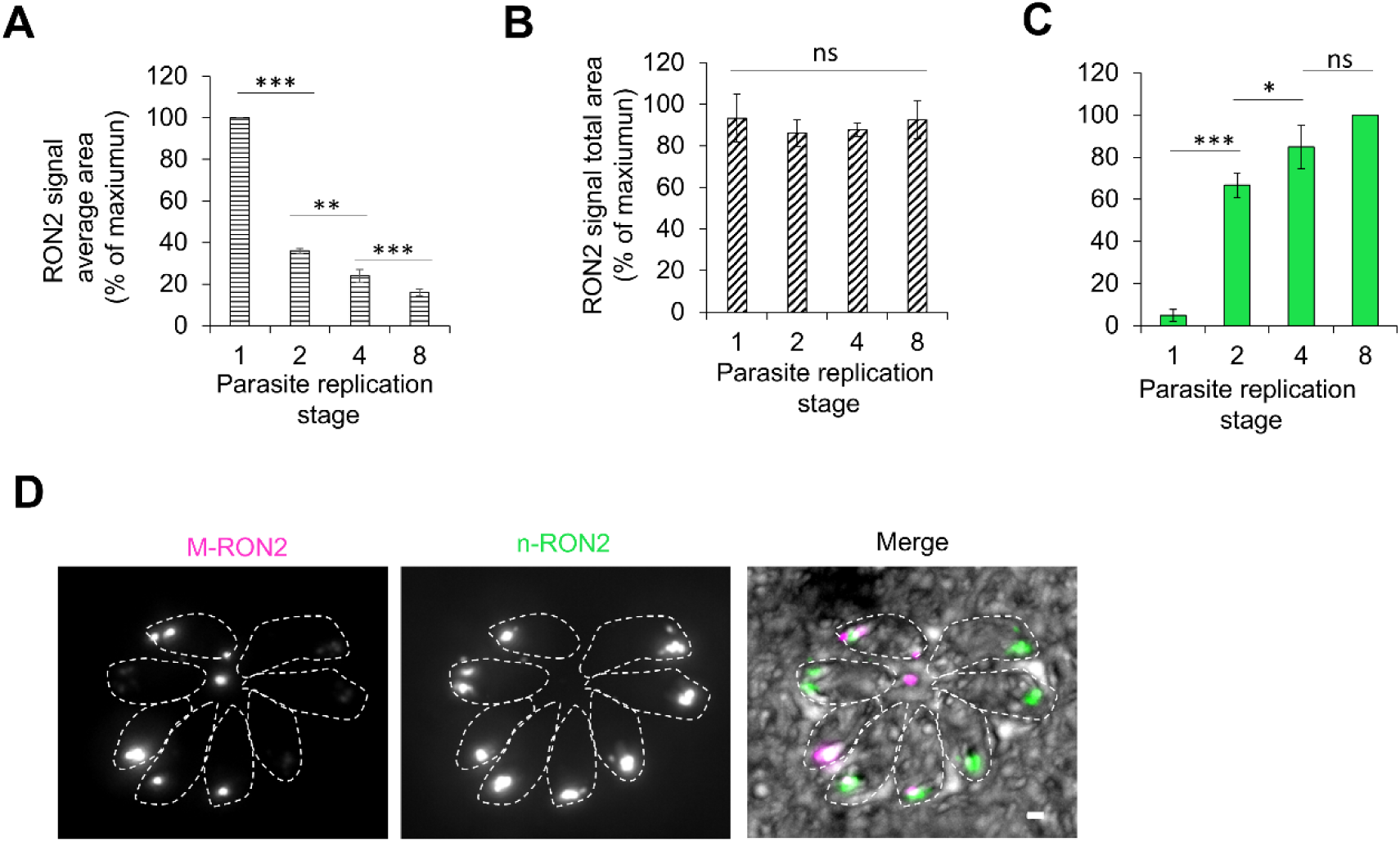
The de novo secretory organelles are generated independently of the maternal. A) Quantification of the average signal area of M-RON2 during replication. The average surface of M-RON2 decrease at each replication suggesting a separation of the mother organelle. B) Quantification of the total signal area of M-RON2 during replication. Despite the decrease of the average signal area, the total surface of M-RON2 remain stable indicating that the totality of the maternal organelles are conserved during replication. C) Fluorescence intensity analysis of n-RON2. D) Representative picture of a vacuole with missing maternal rhoptries in some daughter cell of a single vacuole at stage 8. Magenta: M-RON2, Green: n-RON2. Three biological replicates were used for all analysis; error bars are standard deviations, and the centre measurement of the graph bars is the mean. For each panel A,B,C a total of 289 vacuoles were analysed. All p-values ≤ 0.001 (***), using two-tailed unpaired Student’s t-test. Panels D shows maximum-intensity projections of Z-stack images. All scale bars = 1 µm.

**Figure S5.**
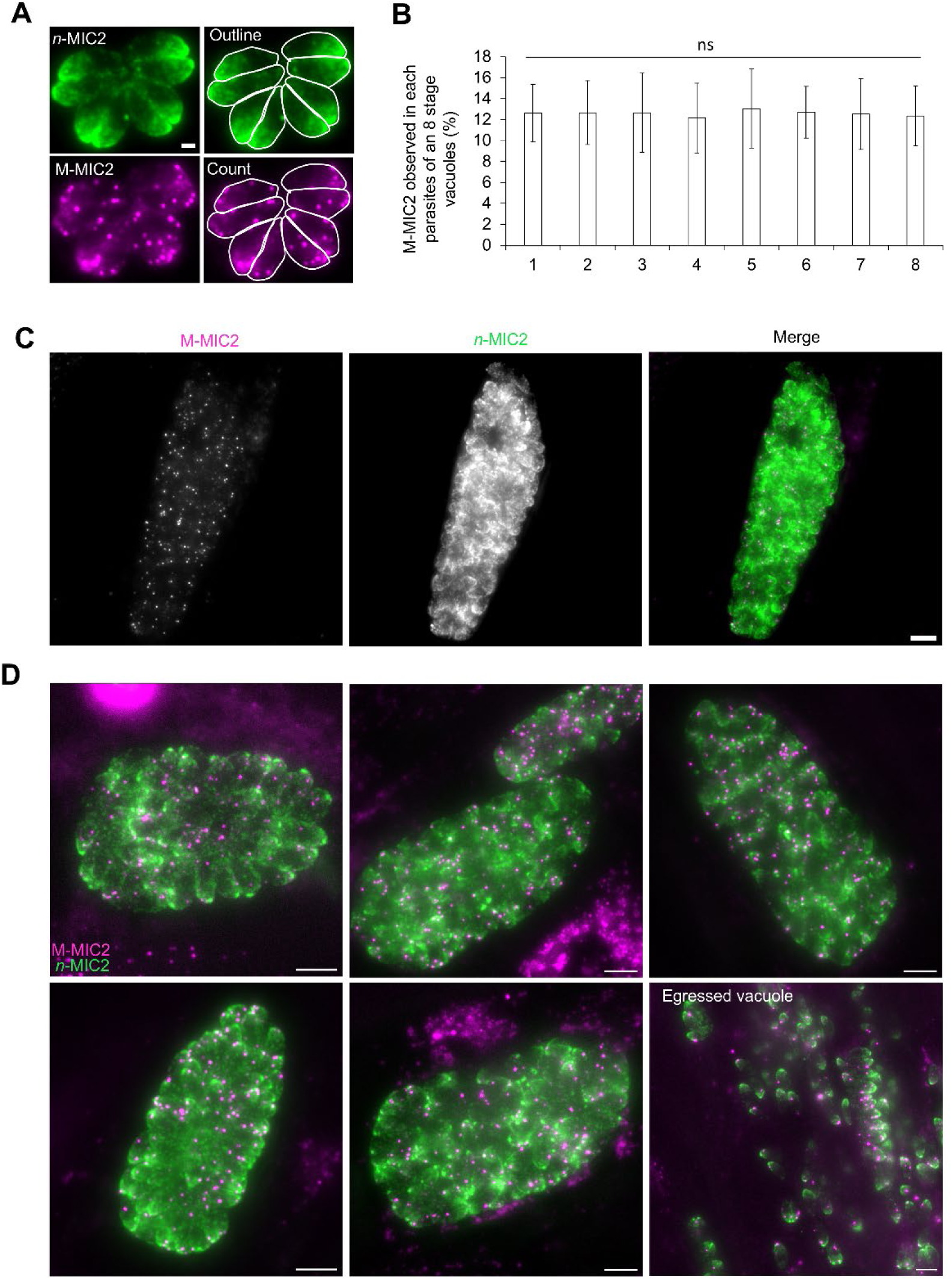
Maternal micronemes are evenly distributed to daughter cells. A) MIC2-Halo parasites at stage 8 were stained for maternal (M-MIC2, magenta) and de novo (n-MIC2, green) MIC2. n-MIC2 signal was used to outline each tachyzoite, numbered 1–8 left to right, and M-MIC2 micronemes were counted per cell. Scale bar = 1 µm. B) Quantification of average M-MIC2 micronemes per tachyzoite shows consistent numbers across daughters, indicating equal distribution. A total of 18 vacuoles were analysed for a total of 1341 isolated M-MIC2 signals. C) Representative image of a late-stage vacuole shows uniform M-MIC2 distribution among all daughter cells. D) Supplementary images from the different replicate that illustrating the inheritance pattern is constant between vacuoles. All p-values ≥ 0.05 (ns), using two-tailed unpaired Student’s t-test. Panels A, C and D show maximum-intensity projections of Z-stack images. Scale bar = 5 µm. Three biological replicates were used; bars show means ± SD. All p-values ≤ 0.001 (***), using two-tailed unpaired Student’s t-test.

**Figure S6.**
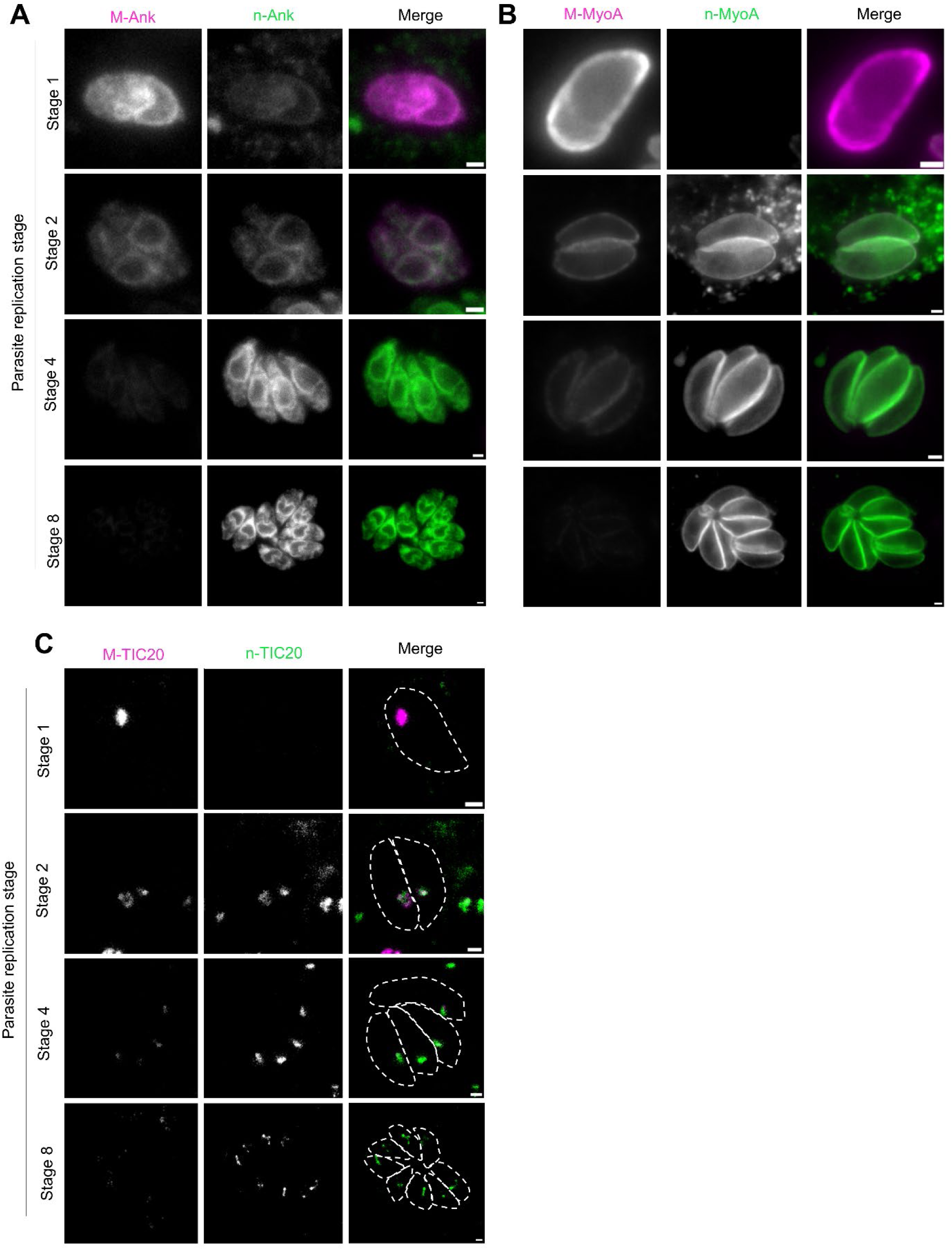
Maternal and *de novo* signal of the other protein of the group 2. All parasite lines were labelled for maternal (pior invasion, Jan 646, magenta) and *de novo* material (after replication, Jan 549, green) A) ANKER1-Halo. Magenta: M-ANKER1, Green: n-ANKER1. B) MyoA-Halo. Magenta: M-MyoA, Green: n-MyoA. C) TIC20-Halo. Magenta: M-TIC20, Green: n-TIC20. Panels displays maximum-intensity projections of Z-stack images. Scale bars = 1 µm.

**Figure S7.**
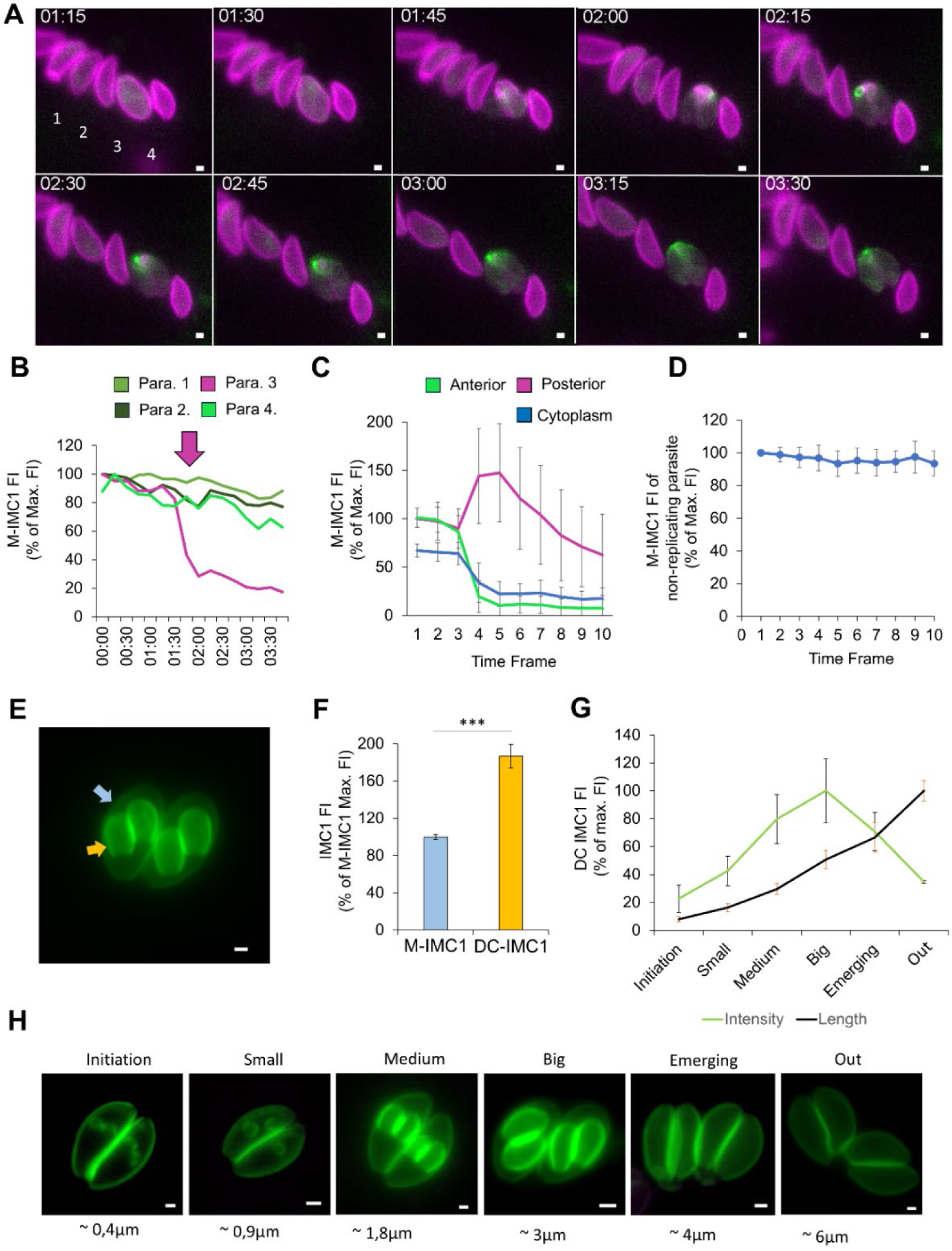
IMC degradation occurs after daughter cell formation but before budding. A) Time-lapse of IMC1-Halo during the first replication cycle, with F-actin visualized via chromobody-emerald. Magenta: M-IMC1, Green: F-actin. M-IMC1 accumulates in the residual body with no redistribution or signal loss in non-replicating parasites. B) Fluorescence intensity tracking of parasites 1–4 (see A) during replication of parasite 3. C) Mean fluorescence intensity across apical, cytoplasmic, and basal regions in 25 replicating parasites. D) Mean M-IMC1 intensity in 25 non-replicating parasites during the same timeframe as C. E) Image showing daughter cells forming within the mother. Green: IMC1-YFP. Blue arrow: mother; yellow arrows: daughters. F) IMC fluorescence comparison between mother and daughters, with mother set to 100%. A total of 47 daughter cells and 75 maternal IMC were analysed. G) IMC intensity in daughters relative to size. Peak intensity occurs when daughters reach ∼3 µm, just before emergence. A total of 100 vacuoles were analysed. H) Representative images of daughter cells at different sizes used for classification. Three biological replicates were used; bars show means ± SD. Panel A displays single optical plane images acquired during live imaging. Panels E and H show maximum-intensity projections of Z-stack images. Scale bars = 1 µm.

**Figure S8.**
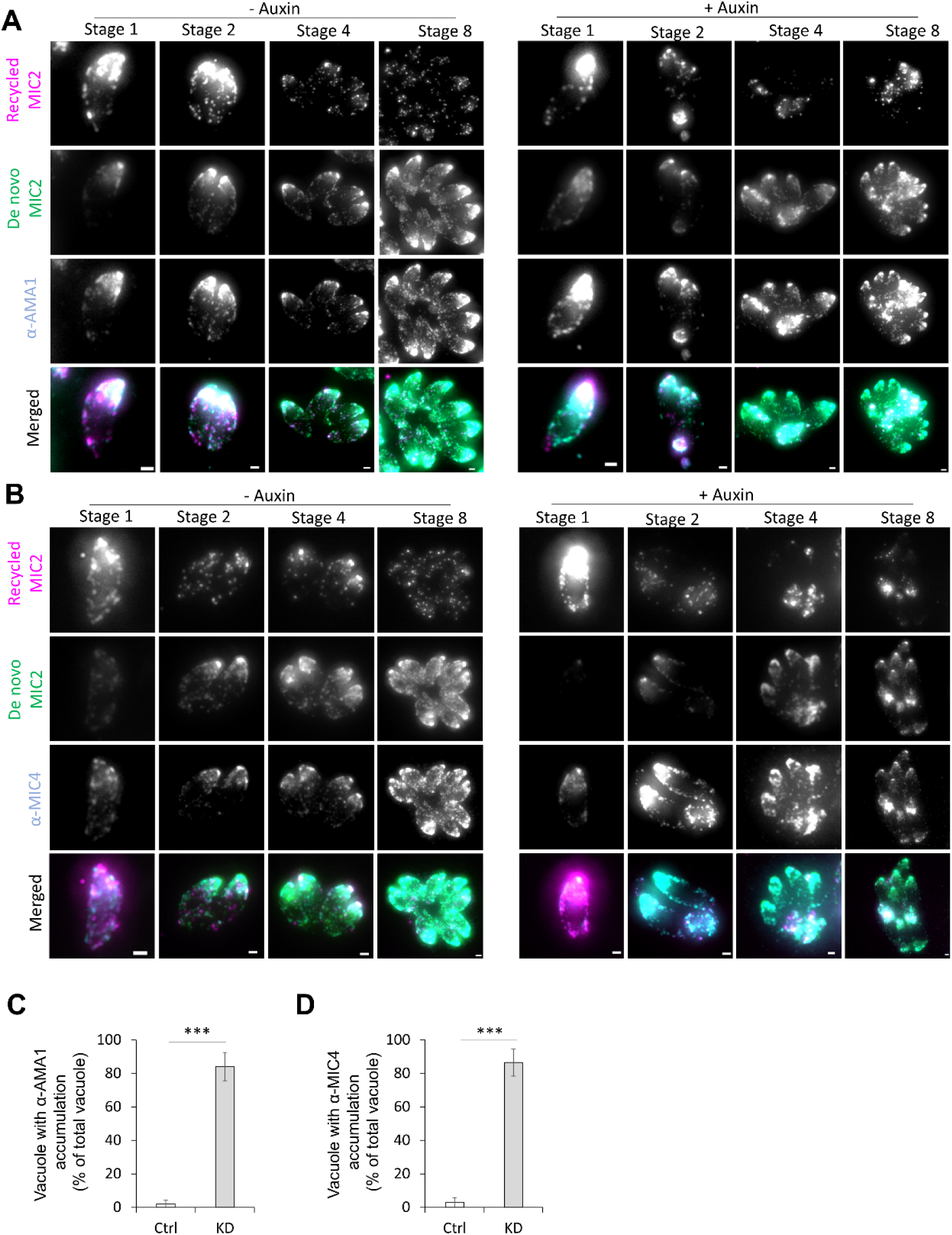
AMA1 and MIC4 recycling is impaired in the absence of MyoF. Effect of MyoF knockdown (KD) on AMA1 inheritance. MyoF-mAID MIC2-Halo parasites were grown ± auxin for 24h. Magenta: M-MIC2, Green: n-MIC2, Cyan: α-AMA1. Without MyoF, AMA1 accumulates in the residual body, mirroring the MIC2-Halo pattern. B) Effect of MyoF KD on MIC4 inheritance. Magenta: M-MIC2, Green: n-MIC2, Cyan: α-MIC4. MIC4 also accumulates in the residual body over time in the absence of MyoF. C-D) Quantification of vacuoles showing α-AMA1 (C) and α-MIC4 (D) accumulation. White: control; Gray: MyoF-KD. Three biological replicates were used; a total of 904 Ctrl and 731 KD vacuoles for AMA1 and 691 Ctrl and 661 KD for MIC4 were analysed. bars show means ± SD. All p-values ≤ 0.001 (***), using two-tailed unpaired Student’s t-test. Panels A and B show maximum-intensity projections of Z-stack images. Scale bars = 1 µm.

**Figure S9.**
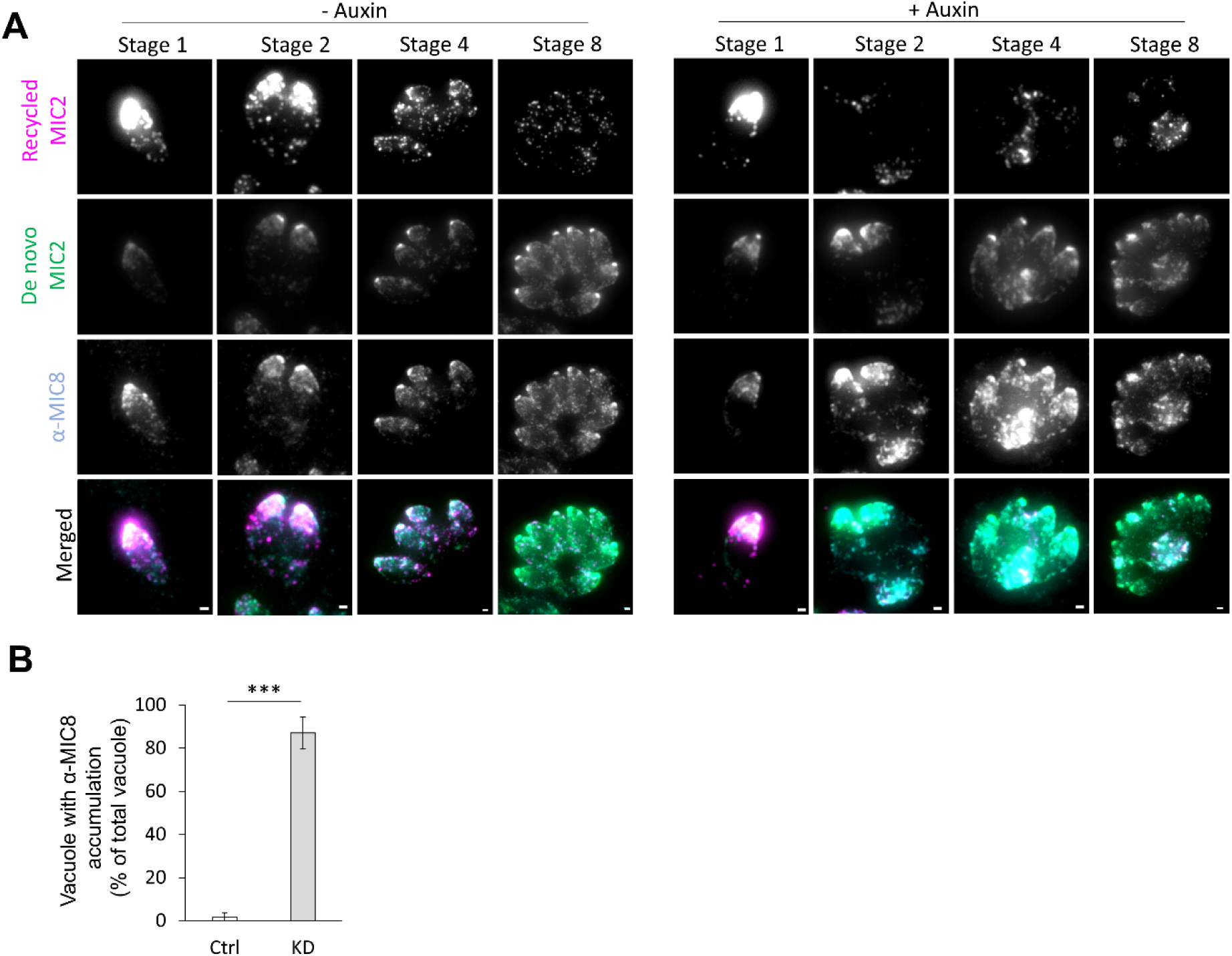
In absence of MyoF, MIC8 is also blocked in its recycling. Impact of MyoF KD on MIC8 inheritance during replication. MyoF-mAID MIC2-Halo parasites were labelled and grown for 24h +/- auxin. MIC8 was visualised using antibodies. Magenta: M-MIC2, Green: n-MIC2, Cyan: α-MIC8. In absence of MyoF, as observed for all the other microneme markers, α-MIC8 accumulate as replication goes on, following a similar patten as MIC2-Halo. **C)** Quantification of the percentage of vacuoles exhibiting accumulation of α-MIC8 in the residual body. White: Control, Gray: MyoF-KD. Three biological replicates were used for all analyses; a total of 900 Ctrl and 671 KD vacuoles were analysed, error bars are standard deviations, and the centre measurement of the graph bars is the mean. All p-values ≤ 0.001 (***), using two-tailed unpaired Student’s t-test. Panels A shows maximum-intensity projections of Z-stack images. All scale bars = 1 µm.

**Figure S10.**
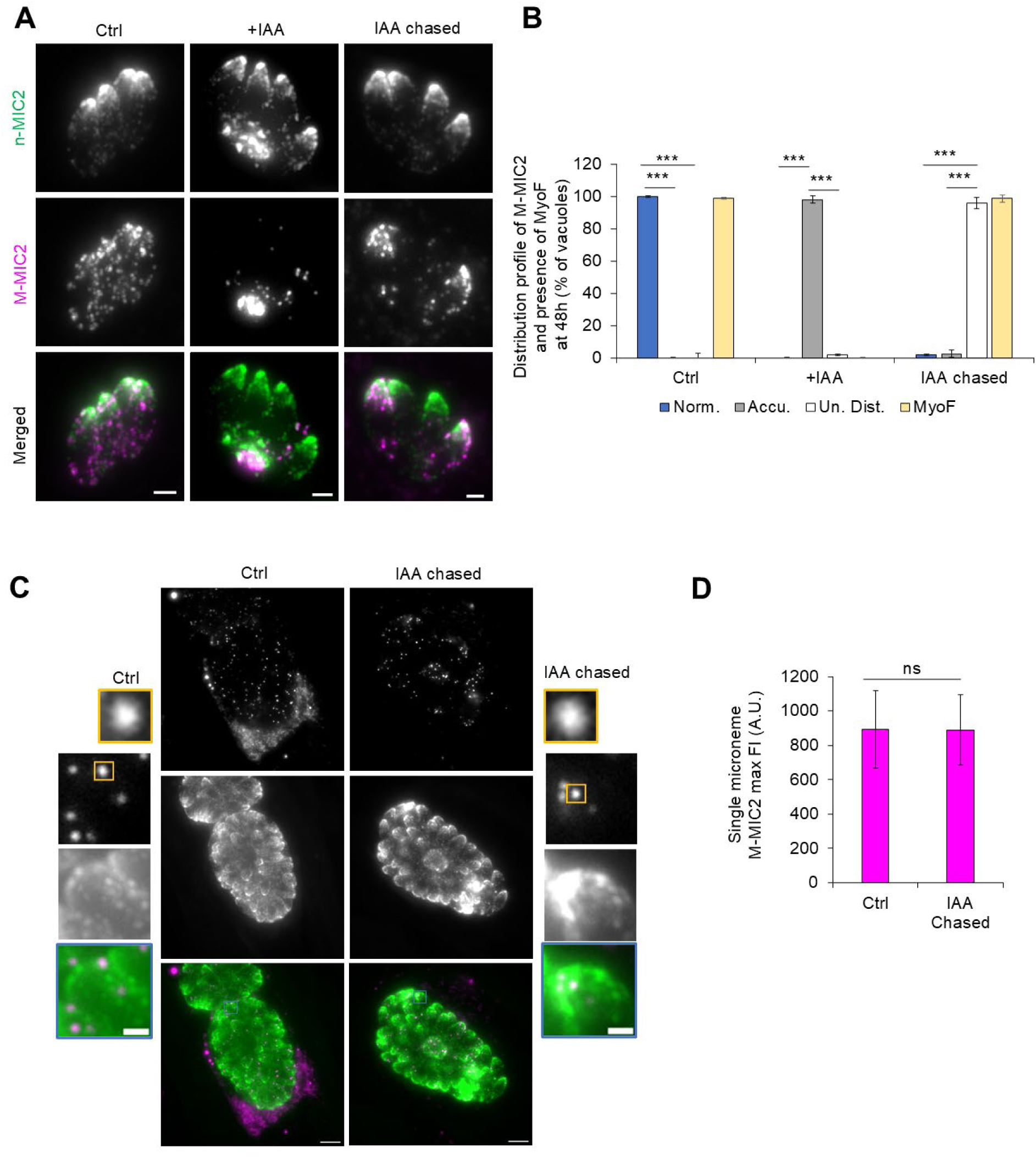
Redistribution of micronemes following auxine washout. A) Representative images of MyoF-mAID MIC2-Halo parasites after 48h in control (no auxin), continuous auxin (Aux), or auxin washout (Aux washed) showing extreme alteration of the redistribution of the M-MIC2. Magenta: M-MIC2, Green: n-MIC2. B) Quantification of vacuoles showing normal, accumulated, or redistributed and MyoF expression. Blue: normal distribution; Gray: accumulated micronemes; White: uneven distribution, Yellow: MyoF. A total of 99 Crl, 120 Kd and 144 auxin washout vacuoles were analysed. C) Representative picture of a control and a vacuole after auxin washout and microneme redistribution, side panel show a focus on an apical part of one tachyzoite of the vacuole and a focus on an isolated microneme signal. D) Quantification of the M-MIC2 fluorescence intensity between control and auxine washout vacuoles. A total of 150 independent micronemes signal were analyzed for both Ctrl and IAA chased. No significant difference was observed suggesting the absence of major degradation process after the 24h sequestration of the micronemes in the residual body. Three biological replicates were used for all analysis. All p-values ≤ 0.001 (***), all p-values ≥ 0.05 (ns) using two-tailed unpaired Student’s t-test. Panels A and C shows maximum-intensity projections of Z-stack images. All scale bars = 1 µm.

**Figure S11.**
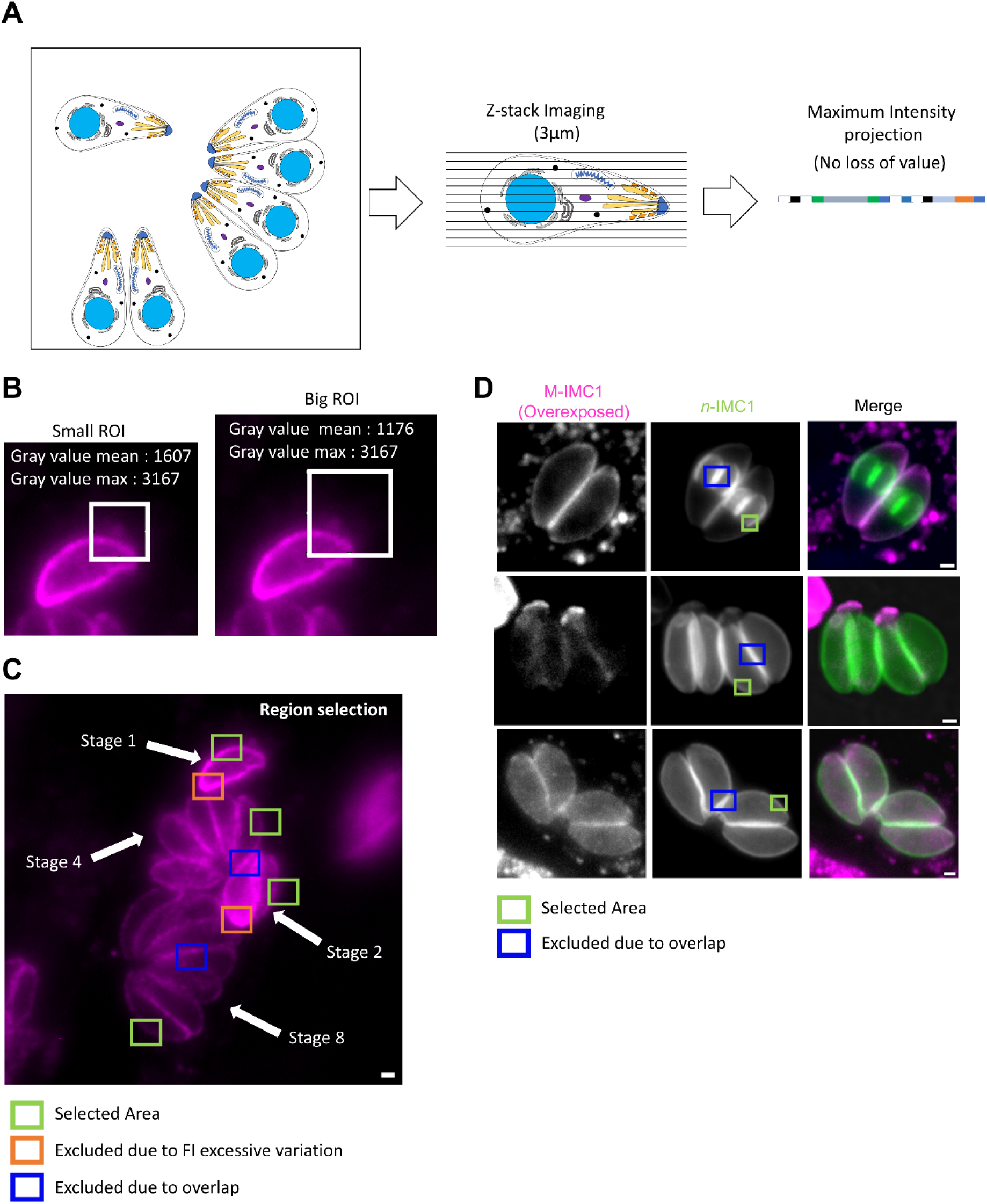
Image processing and fluorescence intensity measurements. Supplementary information supporting the Materials and Methods describing image processing and region selection used for fluorescence intensity quantification throughout the study. (A) Schematic of the imaging strategy, showing Z-stack acquisition centred on the middle of the vacuole, followed by maximum-intensity projection without intermediate image processing. Maximum-intensity projection was chosen over mean projection as it provides a more conservative measure of fluorescence intensity. (B) Illustration of gray-value measurements obtained from the same region of interest (ROI), comparing maximum and mean gray values. While the maximum gray value is independent of ROI size, the mean gray value varies with ROI area. (C) Examples of ROI selection and exclusion for gray-value measurements in parasites at different replication stages.(D) Example of ROI selection for gray-value measurement of mother versus daughter IMC, using the initial maternal IMC signal to discriminate between the two.

**Video S1: Live degradation of the maternal IMC1.**

Magenta: M-IMC1 (Jan646), Green: Cb-Emerald.

**Video S2: Live inheritance of the maternal RON2.**

Magenta: M-RON2 (Jan646), Green: IMC1-YFP.

**Video S3: Live inheritance of the maternal MIC2.**

Magenta: M-MIC2 (Jan646), Green: IMC1-YFP.

**Video S4: Live replication of MyoF mAID MIC2-Halo without auxin**

Magenta: M-MIC2 (Jan646), Green: n-MIC2 (Jan549).

**Video S5: Live replication of MyoF mAID MIC2-Halo with auxin**

Magenta: M-MIC2 (Jan646), Green: n-MIC2 (Jan549).

**Video S6: Live replication of MyoF mAID MIC2-Halo after auxin chase**

Magenta: M-MIC2 (Jan646), Green: n-MIC2 (Jan549).

